# *In vitro* transcription-based biosensing of glycolate for prototyping of a complex enzyme cascade

**DOI:** 10.1101/2024.04.26.591264

**Authors:** Sebastian Barthel, Luca Brenker, Christoph Diehl, Nitin Bohra, Simone Giaveri, Nicole Paczia, Tobias J Erb

**Author notes:** These authors contributed equally to this work (Sebastian Barthel, Luca Brenker). Corresponding authors: Tobias J Erb and Sebastian Barthel.

## Abstract

*In vitro* metabolic systems allow the reconstitution of natural and new-to-nature pathways outside of their cellular context and are of increasing interest in bottom-up synthetic biology, cell-free manufacturing and metabolic engineering. Yet, the prototyping of such *in vitro* networks is very often restricted by time- and cost-intensive analytical methods. To overcome these limitations, we sought to develop an *in vitro* transcription (IVT)-based biosensing workflow that offers fast results at low-cost, minimal volumes and high-throughput. As a proof-of-concept, we present an IVT biosensor for the so-called CETCH cycle, a complex *in vitro* metabolic system that converts CO_2_ into glycolate. To quantify glycolate production, we constructed a sensor module that is based on the glycolate repressor GlcR from *Paracoccus denitrificans*, and established an IVT biosensing off-line workflow that allows to measure glycolate from CETCH samples from the µM to mM range. We characterized the influence of different cofactors on IVT output and further optimized our IVT biosensor against varying sample conditions. We show that availability of free Mg^2+^ is a critical factor in IVT biosensing and that IVT output is heavily influenced by ATP, NADPH and other phosphorylated metabolites frequently used in *in vitro* systems. Our final biosensor is highly robust and shows an excellent correlation between IVT output and classical LC-MS quantification, but notably at ∼10-fold lowered cost and ∼10 times faster turnover time. Our results demonstrate the potential of IVT-based biosensor systems to break current limitations in biological design-build-test cycles for the prototyping of individual enzymes, complex reaction cascades and *in vitro* metabolic networks.

## Introduction

Synthetic biochemistry aims to reconstruct biological functions outside of a living cell (“cell-free”). Prominent examples are efforts to reconstitute natural (or new-to-nature) pathways from purified enzymes *in vitro*^1–8^. Such approaches allow studying the fundamental design principles and function of metabolic networks, but also bear great application potential. For example, recent work demonstrated the cell-free conversion of the greenhouse gas carbon dioxide (CO_2_) into polyketides, terpenes and antibiotic precursors or the valorization of glucose and other low-cost precursors into monoterpenes and cannabinoids.

Compared to *in vivo* systems, cell-free metabolic networks are highly flexible in their composition, can be precisely modified and customized, and allow biochemical reactions to take place in “non-physiological” conditions^9^. Through cell-free systems, a rapid optimization of reaction compositions is possible without the need for molecular cloning, which minimizes time and cost. Consequently, lysate-based cell-free systems have been increasingly used to prototype pathways for the optimal combination and stoichiometry of individual enzymes and components^10–14^. In several cases, these optimized *in vitro* pathways could be also successfully implemented *in vivo*. Altogether, this showcases the capabilities of cell-free systems as a broad tool for *in vitro* and *in vivo* metabolic engineering purposes.

To optimize complex biological systems with minimal experimental effort, Pandi and coworkers recently reported a versatile workflow (METIS) that combines laboratory automation with active learning to explore the combinatorial space in iterative design-build-test cycles for (local) optima^15^. METIS successfully helped to improve several biological systems^3,16–18^, including an *in vitro* CO_2_ fixation cycle of 27 different variables (CETCH cycle^2,19^). The CETCH cycle converts CO_2_ into glycolate and could be improved by more than 10-fold through METIS. Although active learning-guided workflows are able to drastically minimize the number of samples screened, the screening phase still heavily relies on the use of costly and time-intensive instrumental analytics. In the case of the CETCH cycle, >3,000 samples were analyzed by liquid chromatography-mass spectrometry (LC-MS) which requires 12 min per sample for glycolate quantification at a cost of approximately US$7 (Supplementary Note 1). We therefore set out to explore a low-cost and well-scalable *in vitro* transcription (IVT)-based biosensing method to increase the throughput of screening campaigns in complex conditions.

To establish such an IVT biosensing method, we turned our attention to a system called RNA Output Sensors Activated by Ligand Induction (ROSALIND), which was recently developed to detect pollutants in water samples^20^. The ROSALIND system consists of a linear DNA template encoding the sequence of an RNA aptamer (“Three-way Junction dimeric Broccoli”, *3WJdB*)^21^ (Figure 1A). The *3WJdB* aptamer is expressed under the control of a T7 promoter and an operator sequence that is repressed by an allosteric transcription factor (aTF). Only in the presence of its specific effector, the aTF releases the operator sequence to allow *3WJdB* expression, which in turn results in a green fluorescent readout by stabilizing the fluorogenic dye DFHBI-1T in its fluorescent state.

**Figure 1:**
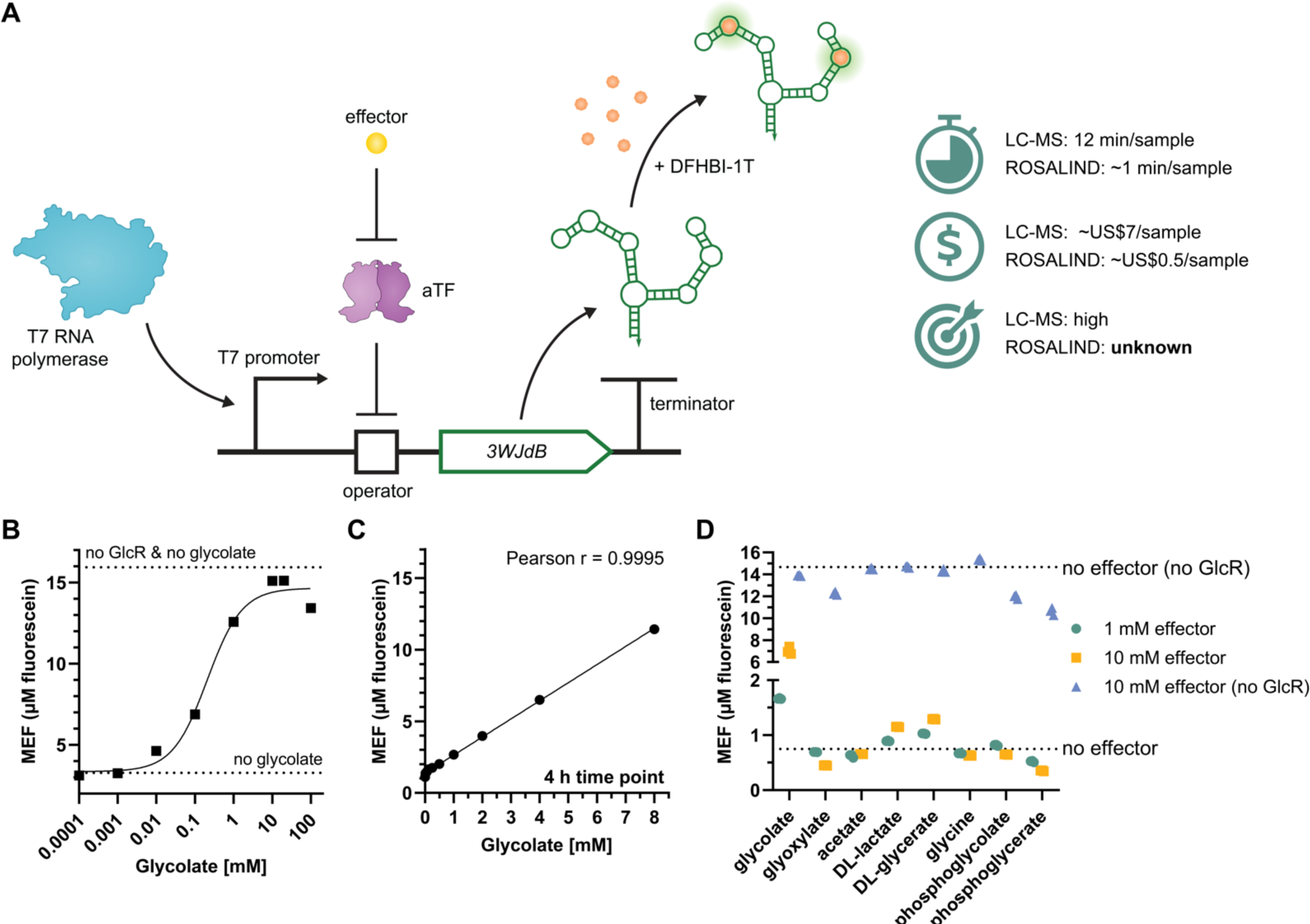
Characterization of a glycolate-responsive *in vitro* transcription-based biosensor module. **A:** The ROSALIND system is based on the controlled expression of the *3WJdB* RNA aptamer and the correlating fluorescence signal of the *3WJdB*:DFHBI-1T complex. Expression is regulated by a transcriptional repressor that binds to an operator sequence downstream of a T7 promoter. The repression is lifted in a dose-responsive manner by binding of an effector molecule to the aTF. This system allows faster and cheaper sample measurement in microtiter plates than analysis by LC-MS, but its precision is yet unknown. Time estimates refer to glycolate quantification. **B, C:** The dose-response curve of the GlcR sensor module to 100 nM to 100 mM glycolate (16 h time point) shows an operational range from 10 µM to 20 mM glycolate with an excellent correlation between 16 µM and 8 mM glycolate and a 10-fold dynamic range. Note that IVT biosensing reactions are highly time-sensitive (Supplementary Figure 5). **D:** Promiscuity assay of GlcR shows a low-level dose response to DL-lactate and DL-glycerate. Raw fluorescence data are standardized to MEF (µM fluorescein). Data are the mean of *n*=3 technical replicates ± s.d. IVT output without effector molecule and without MGlcR is shown as horizontal dotted lines.

Here, we demonstrate a ROSALIND-based biosensing workflow for the rapid prototyping and screening of complex *in vitro* metabolic systems, using the CETCH cycle as proof-of-principle. We developed a glycolate-responsive sensor module to read out the glycolate-forming activity of the CETCH cycle, and investigated the inhibitory effects of CETCH cycle components on the IVT system. We identified the availability of free Mg^2+^ as a critical factor for establishing highly robust and quantitative sensing of glycolate production. Notably, the IVT-based biosensing workflow reduces screening costs by an order of magnitude and reduces the analysis time of large sample sets from several days to approximately eight hours. This work not only demonstrated that IVT-based reporter systems are suitable for screening complex *in vitro* systems, but also identified critical components and bottlenecks in setting up robust IVT-based screens under complex and challenging conditions. Our work paves the way for the development of similar IVT-based reporter systems and further guides ongoing efforts to integrate *in vitro* metabolic networks and *in vitro* transcription-translation systems^22^ toward constructing a synthetic cell^23,24^.

## Results

### Establishing GlcR as glycolate-responsive IVT-based biosensor module

To develop IVT-based reporter systems for *in vitro* metabolic networks, we chose the CETCH cycle as example, because of its pioneering role in synthetic CO_2_ fixation, its biological complexity (involving a total of 17 enzymes), as well as its recent use in a METIS-assisted optimization workflow in which 1,000 different combinations (3,000 samples) had been already tested^15^.

The end product of the CETCH cycle is glycolate, which is produced from CO_2_. We therefore set out to construct a glycolate-responsive sensor module from a ROSALIND DNA template and a suitable allosteric transcription factor (aTF). A glycolate-responsive transcription factor, GlcC from *Escherichia coli,* was previously described. However, this protein acts as a transcriptional activator^25–27^, which made it incompatible with the T7 promoter-based IVT system of ROSALIND, which strictly relies on transcriptional repression. Thus, we turned our attention to another aTF from *Paracoccus denitrificans* that was recently reported to regulate glycolate assimilation in the β-hydroxyaspartate cycle (BHAC)^28^. This GntR family transcriptional repressor, named GlcR, is encoded by *pden4400,* binds the intergenic sequence *pden4399-4400* and was shown to unbind in the presence of glycolate^29^, which made the protein an interesting candidate for our envisioned IVT-based biosensor.

We purified GlcR as a fusion protein with N-terminal maltose-binding protein (MGlcR) and confirmed its binding and unbinding from the intergenic sequence between *pden4400* and *pden4397-4399* in the absence and presence of glycolate, respectively, by electrophoretic mobility shift assays (EMSA) (Supplementary Figure 1A). To identify fragments carrying a putative operator site (*glcO*, Supplementary Figure 1B), we next split the 150 base pair (bp)-long intergenic sequence into six fragments, with each fragment composed of ∼60 bp in length and ∼30 bp overlap with neighboring fragments. The sixth fragment also encoded the first 51 bp of *pden4399*. EMSA showed that MGlcR bound to four of the six fragments (fragments #2-5), and in particular to fragment #3, which was bound by MGclR ∼2 to 4-fold stronger as fragments #2, #4, and #5, indicating that GlcR has multiple operator sites.

We then focused on the putative operator site in fragment #3. However, further splitting of fragment #3 into 20 bp and 30 bp-long fragments completely abolished MGlcR binding, suggesting that the operator site spans more than 30 bp (Supplementary Figure 1C). We next removed base pairs in 4 bp steps from the 5’ end of fragment #3 and prepared eight ROSALIND templates encoding putative *glcO* sequences between 60 and 32 bp length (named according to their length, i.e., the 60 bp-long sequence was named *glcO_60_*) as part of a P_T7_-*glcO*-*3WJdB* expression cassette (Supplementary Note 2). When tested for *3WJdB* expression in the presence or absence of MGlcR, all eight constructs showed repression between 3-fold (*glcO_48_* and *glcO_52_*), and up to 8-fold and 16-fold in the case of *glcO_60_* and *glcO_36_*, respectively (Supplementary Figure 2A).

We continued with the two best sensor constructs, *glcO_36_* and *glcO_60_*, and titrated MGlcR over a constant DNA template concentration (25 nM). At 1.25 µM MGlcR (i.e., 50x aTF:DNA template ratio) *glcO_36_* showed a ∼80-fold repression, while *glcO_60_* required 5 µM MGlcR (i.e., 200x aTF:DNA template ratio) to reach a similar level of repression (Supplementary Figure 2B). At these concentrations, both constructs showed a ∼3-fold de-repression with 10 mM glycolate (Supplementary Figure 2C). We selected *glcO_36_* as final construct and sought to further increase sensitivity of the system by reducing the total aTF concentration. It has recently been shown that removing excess aTF molecules, which act as effector chelators, improves the sensitivity of ROSALIND sensor modules^20^. When lowering the DNA template to 15 nM or 5 nM DNA (and keeping the aTF:DNA ratio at 50x (i.e., 0.75 µM and 0.25 µM MGlcR)), glycolate-induced de-repression increased by 6-fold and 10-fold, respectively, while the signal was only reduced by 8% and 50%, respectively, resulting in a good balance between glycolate sensitivity and total output signal (Supplementary Figure 2D). We tested the influence of the T7 RNA polymerase (RNAP) preparation onto the signal (Supplementary Figure 3) and confirmed that the sensor was functional in HEPES buffer between pH 7.2 and 7.8, the buffer conditions of the CETCH cycle (Supplementary Figure 4). As standard conditions for all subsequent IVT-based biosensing experiments, we chose HEPES buffer pH 7.8 with 15 nM *glcO_36_*, 750 µM MGlcR, and in-house T7 RNAP, which we refer to as the GlcR sensor module.

### The GlcR sensor module is operational over three orders of magnitude

To determine the operational range of the GlcR sensor module, we tested the response of the sensor to glycolate concentrations over six orders of magnitude (from 100 nM to 100 mM), which defined a limit of detection at 10 µM glycolate and showed inhibition at glycolate concentrations above 20 mM (Figure 1B, Supplementary Figure 5A). The GlcR sensor module showed an excellent linear response between 16 µM and 8 mM (Pearson r = 1.0) and a dynamic range of 10.3-fold after 4 h of incubation (Figure 1C, Supplementary Figure 5C-D).

We also tested the specificity of the GlcR sensor module with seven structurally- and context-related small organic acids: glyoxylate, acetate, DL-lactate, DL-glycerate, glycine, 2-phosphoglycolate, 3-phospho-D-glycerate. Notably, the sensor was highly specific for glycolate and showed no response with other C2 acids, including glyoxylate and the amino acid glycine. However, we observed some dose-responsive signal with DL-lactate and DL-glycerate, indicating some promiscuity of GlcR with C3 alpha-hydroxy acids (Figure 1D, Supplementary Figure 6). However, since these C3 acids are not part of the CETCH cycle, we concluded that the GlcR sensor module could be used for the envisioned IVT-based biosensing system.

### Non-enzyme components of CETCH influence on-line IVT analysis

To assess whether on-line (i.e., direct) biosensing of CETCH samples would be possible with our IVT biosensor, we next tested the compatibility of IVT with components of the CETCH cycle. The CETCH cycle consists of 27 components – 17 enzymes and ten non-enzyme components, which include substrates (bicarbonate, propionyl-CoA), cofactors (ATP, MgCl_2_, NADPH, coenzyme A (CoA), coenzyme B12), metabolites for energy supply (formate, creatine phosphate), and a buffer reagent (HEPES) (Figure 2A). In addition to these ten non-enzyme components of the cycle, all CETCH enzymes are kept in 20% glycerol, propionyl-CoA oxidase (Pco) and methylsuccinyl-CoA oxidase (Mco) are stored additionally with flavin adenine dinucleotide (FAD), and commercially available enzymes such as T7 RNA polymerase with β-mercaptoethanol (β-ME).

**Figure 2:**
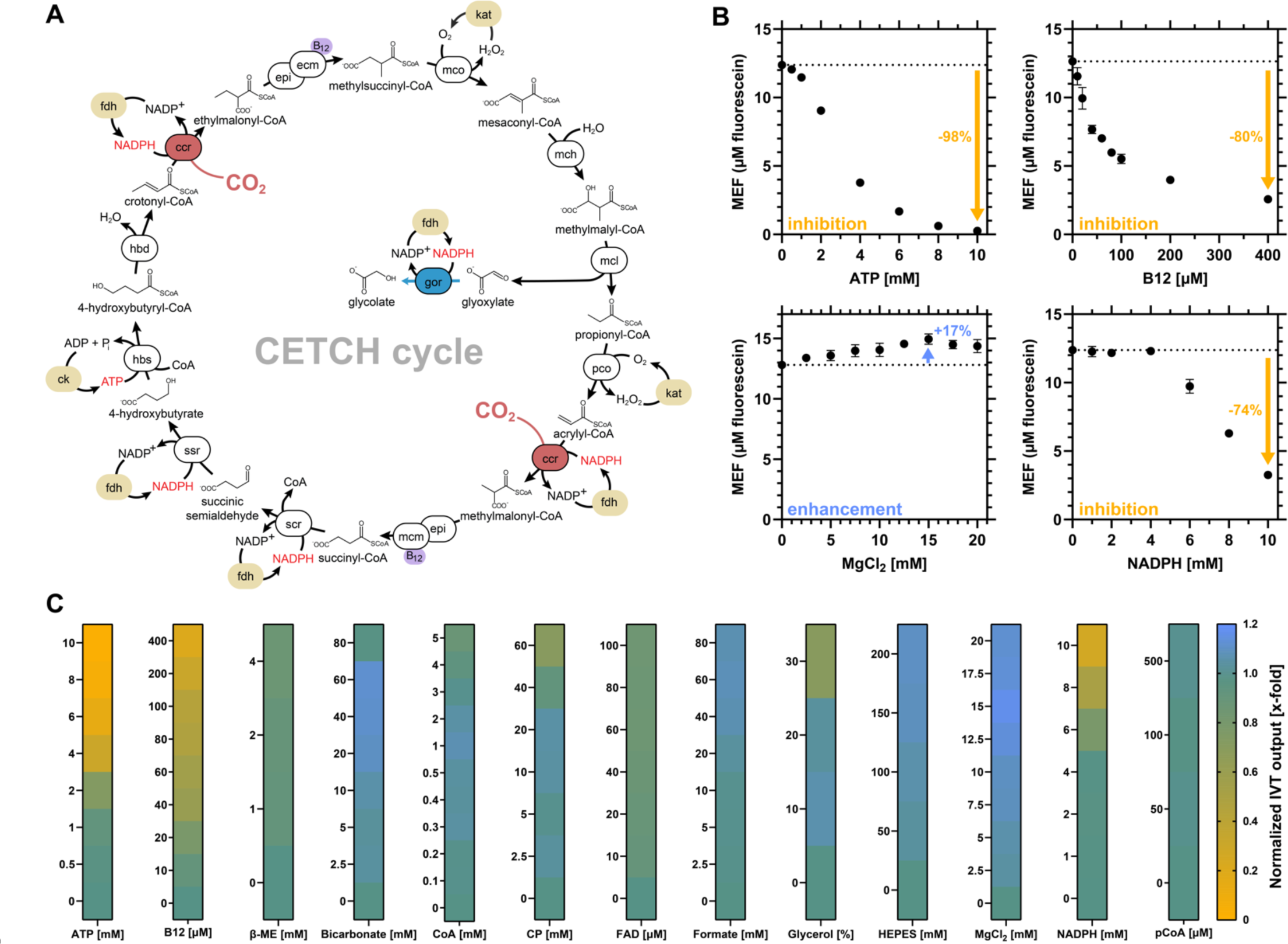
Influence of CETCH cycle components on *in vitro* transcription in the absence of GlcR. **A:** Reaction sequence of the CETCH cycle to convert CO_2_ into glycolate^2,19^. 10 non-enzymatic components are actively involved in the cycle, 3 additional components are required to maintain enzyme activity during storage. **B:** Titration of ATP, B12, MgCl_2_ and NADPH concentrations in IVT reactions shows a dose-dependent influence of each component on IVT output after 4 h. Detailed data for 9 additional components are shown in Supplementary Figure 7. Raw fluorescence data are standardized to MEF (µM fluorescein). Data are the mean of *n*=3 technical replicates ± s.d. **C:** Heatmap describing the influence of all 13 non-enzyme components of the CETCH cycle on IVT output (as shown in **B** and Supplementary Figure 7). Data are normalized to IVT output in the absence of the respective component. Orange, blue and green colors indicate inhibition, enhancement and no effect, respectively, of the screened component at the indicated concentration.

We investigated the individual influence of these 13 different non-enzyme components onto our basic IVT system (without the GlcR sensor module). For the ten non-enzyme components of the CETCH cycle, we tested concentrations in our IVT that were previously used during METIS-assisted optimization of the cycle by Pandi and coworkers^15^. For β-ME, FAD, and glycerol, we sampled a wider range of concentrations (Supplementary Table 5). Notably, 8 of the 13 non-enzyme components inhibited the IVT reaction, with ATP, NADPH, and B12 showing strong inhibition of up to 74-98% at high concentrations (Figure 2B-C, Supplementary Figure 7). ATP inhibition was partially due to an increased competition between ATP and the other three rNTPs (Supplementary Figure 8). However, the majority of ATP inhibition seemed to be caused by chelation of free magnesium ions, which are essential for T7 RNA polymerase activity, similar to the inhibition of phi29 DNA polymerase by rNTPs^30^. We further speculated that NADPH, creatine phosphate (CP), CoA and FAD, followed a similar inhibitory mechanism. High concentrations of glycerol inhibited by 30%. CoA, FAD and β-ME showed minor inhibition between 12 and 15%, while propionyl-CoA showed no effect. In contrast, four non-enzyme components, bicarbonate, MgCl_2_, formate and HEPES increased IVT output by up to 15% (Figure 2B-C, Supplementary Figure 7).

Overall, the complex (and partially adverse) effects of the different non-enzyme components onto IVT showed that on-line measurements cannot be simply used for IVT-based biosensing in complex, varying conditions of the CETCH cycle. We thus decided to work with an off-line workflow, in which samples are quenched and diluted 1:10 before analysis to minimize the effects of the non-enzyme components onto our IVT biosensor.

### Establishing off-line IVT sensing of CETCH samples with one enzyme component varied

We next developed an off-line biosensing workflow, in which CETCH cycle variants are run first, and their output is analyzed by our IVT biosensor in a subsequent step (Figure 3A). To quench CETCH reactions, we separated small molecules from enzymes by filtration through a 10 kDa molecular weight cutoff (MWCO) membrane before analysis of the filtrate (Supplementary Figure 9) in a 1:10 dilution.

**Figure 3:**
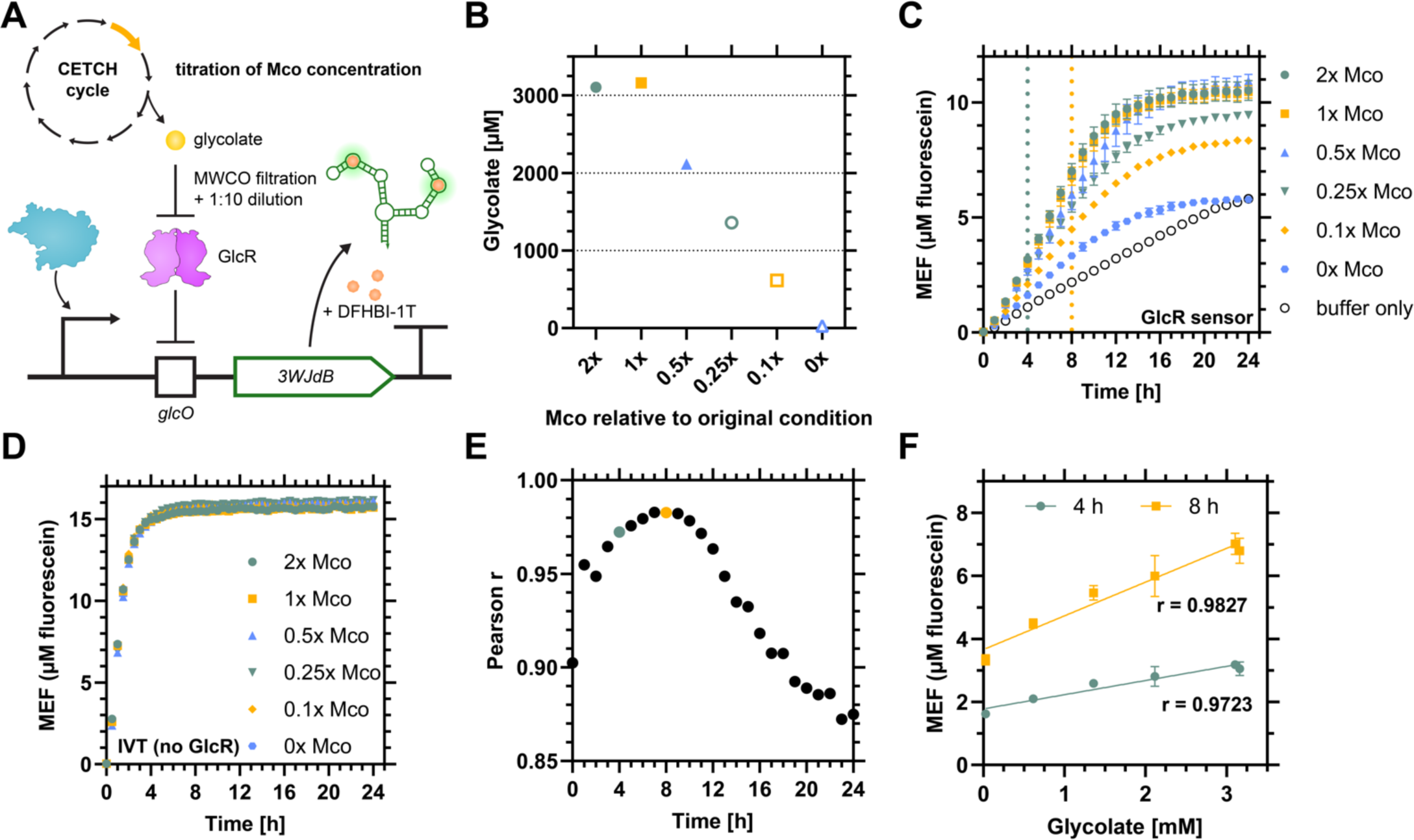
Glycolate sensing from CETCH cycle samples with a single component, the concentration of enzyme Mco, varied. **A:** Schematic of the experimental setup. **B:** LC-MS quantification of glycolate from six CETCH cycle samples with titrated Mco concentration measured in technical triplicates. **C:** Time course of glycolate measurement using the GlcR sensor module. Time points shown in **F** are indicated as vertical dotted lines. **D:** Time course of IVT measurement in the absence of GlcR showing no differences in inhibition by CETCH cycle samples. **E, F:** Correlation between GlcR sensor module output and LC-MS quantification (as shown in **B**) over time. The quality of the correlation is time-sensitive and worsens as soon as the first IVT reactions plateau. Data of 4 h and 8 h time points (indicated in green and orange, respectively) are exemplarily shown in **F.** Raw fluorescence data are standardized to MEF (µM fluorescein). Data are the mean of *n*=3 technical replicates ± s.d. (**C, D, F**).

As a proof-of-concept, we measured glycolate production from quenched CETCH samples, in which only one enzyme was varied. To that end, we titrated methylsuccinyl-CoA oxidase (Mco), a critical enzyme known to limit the productivity of the CETCH cycle^15^. We ran six CETCH cycle reactions (day 7, condition 15 of Pandi et al.^15^, Supplementary Table 6), with different Mco concentrations (0-52 µM). These six conditions yielded different glycolate concentrations (Figure 3B), which our off-line IVT biosensor was able to quantify with high correlations. The time course showed excellent correlation at 4 h (r = 0.97) and between 7 and 9 h (r >0.98), after which the IVT reactions started to plateau and other factors became limiting (Figure 3C,E-F). Overall, these results demonstrated that our off-line IVT-based biosensor workflow is able to screen the productivity of CETCH cycle variants with varying enzyme concentrations.

### Optimizing off-line sensing of CETCH samples for multiple components varied

We next tested whether our off-line IVT biosensor workflow was able to quantify glycolate concentrations from CETCH samples of highly diverse composition (enzymes and non-enzyme components varied, Figure 4A). We prepared six CETCH samples with known productivity (day 7 of Pandi et al., as specified in Supplementary Table 6) and compared their IVT readout with LC-MS-based quantification of glycolate (Figure 4B). Overall, the GlcR module and LC-MS-based method showed a r = 0.77 after 4 h (Figure 4C,D, Supplementary Figure 10), with sample 2 underestimating, and sample 3 overestimating the actual glycolate concentrations, respectively.

**Figure 4:**
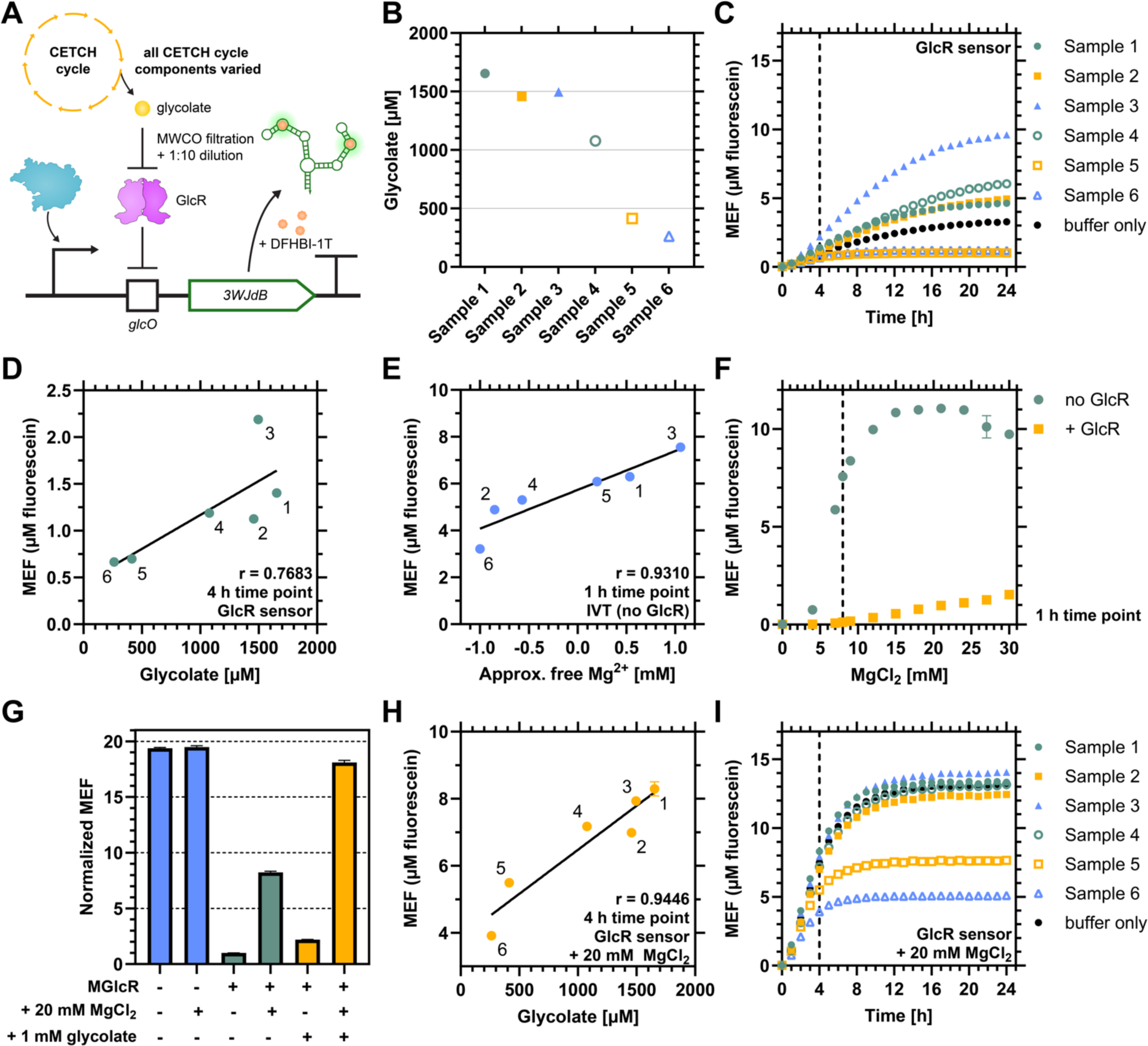
Glycolate sensing from CETCH cycle samples with varied concentrations of non-enzyme components and enzymes. **A:** Schematic of the experimental setup. **B:** LC-MS quantification of glycolate from six CETCH cycle samples of different compositions, measured in technical triplicates. **C, I:** Time course of glycolate measurement with the GlcR module without and with the addition of 20 mM MgCl_2_, respectively. **D, H:** Correlation between GlcR module output (4 h time point, indicated as dashed lines in **C, I**) and LC-MS quantification (as shown in **B**). See Supplementary Figure 10B for correlation coefficients of hourly time points. **E:** Correlation between free Mg^2+^ and IVT output in the absence of GlcR. Free Mg^2+^ is approximated as the concentration difference of added MgCl_2_ and Mg^2+^-binding to estimate the change in free Mg^2+^ upon addition of the CETCH sample to the GlcR sensor module. **F:** Titration from 0 to 30 mM MgCl_2_ showed a dose-dependent IVT output in the presence and absence of GlcR. In the presence of 750 nM GlcR, the leakiness of the repressed module increased linearly, whereas in the absence of GlcR, the IVT output was bell-shaped. **G:** Effect of additional 20 mM MgCl_2_ on the constitutive (blue), repressed (green) and de-repressed (orange) state of the GlcR sensor module. Elevated MgCl_2_ concentrations increased the leakiness of the module but did not affect the response to 1 mM glycolate. Data shown are 4 h time points, normalized to data with only MGlcR added. **C-I:** Raw fluorescence data are standardized to MEF (µM fluorescein). Data are the mean of *n*=3 technical replicates ± s.d.

Strikingly, in both samples, the concentration of (free) MgCl_2_ seemed to be the critical factor. Sample 3 contained the highest MgCl_2_ concentration (17.5 mM) and low concentrations of Mg^2+^-binding cofactors (6.95 mM ATP, NADPH, coenzyme A). In contrast, sample 2 contained the lowest MgCl_2_ concentration (2.5 mM) and high amounts of Mg^2+^-binding cofactors (11 mM ATP, NADPH, coenzyme A). Because Mg^2+^ is the cofactor of T7 RNA polymerase and its availability is essential for IVT (see above), we speculated that Mg^2+^ availability was strongly affecting the read-out in these samples. This was supported by the fact that when we measured the effect of the six CETCH samples on the IVT system without the GlcR module, IVT output correlated well with the approximated concentration of free Mg^2+^ (Figure 4E, r = 0.93).

We therefore decided to increase the overall Mg^2+^ concentration in our biosensor system to minimize the effect of CETCH cycle samples onto Mg^2+^ availability. We examined the Mg^2+^ dependence of the IVT system and the GlcR module in the range of 0 to 30 mM MgCl_2_ (Figure 4F). Between 15 and 25 mM MgCl_2_, the de-repressed IVT system showed a broad plateau, while the Mg^2+^ effect on the repression of the system by GlcR was relatively small, indicating that this MgCl_2_ concentration range was useful for robust sensing.

Indeed, when increasing the MgCl_2_ concentrations from 8 mM by 20 mM in our IVT-based system, this significantly improved the correlation between GlcR module output and LC-MS to r = 0.94 (Figure 4H, Supplementary Figure 10), albeit at some increase of total output signal (Figure 4I), caused by a higher baseline expression of the system (Figure 4G). Overall, however, this setup established our GlcR IVT biosensor as reliable glycolate quantification system that worked robustly across different conditions. This was further confirmed by probing *E. coli* lysate spiked with glycolate (Supplementary Figure 11), demonstrating the possibility to use our IVT-based biosensing also in bacterial lysates that have become an important platform for pathway prototyping, recently^10,11,13,31^.

## Discussion

Here, we explored the potential of IVT-based biosensors in cell-free manufacturing, and in particular synthetic biochemistry. As a proof-of-concept, we tested the sensing of glycolate production from the CETCH cycle, a synthetic CO_2_ fixation cycle, which was only possible by LC-MS analysis, thus far. Our experiments demonstrate that IVT-based biosensing of complex samples is feasible, with excellent and robust correlations.

Key to establish our biosensor was the finding that IVT-based sensing is highly sensitive to components that are commonly used in *in vitro* systems, including ATP, NADPH, and other nucleotide-based and phosphorylated cofactors, which strongly inhibit IVT activity. These results are in line with recent studies that investigated the inhibition of reconstituted transcription, translation and DNA replication systems^30^. In our study, we extend these findings by providing a fitness landscape of T7 RNA polymerase-based transcription in the presence of common metabolic cofactors. This data is not only relevant for efforts to establish other IVT-based biosensors, but might be also helpful for efforts of integrating metabolic and *in vitro* transcription-translation systems towards constructing a synthetic cell^22–24^, and to prototype genetic (RNA) circuits for cell-free biosensing and biocomputing under complex conditions and within artificial compartments^32–36^.

On a more practical note, our work shows that IVT biosensing offers several advantages over classical metabolic quantification methods, because it allows to improve throughput and cost efficiency, and does not rely on expansive analytical instrumentation. For example, during METIS-assisted optimization of the CETCH cycle, Pandi and coworkers screened glycolate production in a 384-well format with LC-MS^15^. At a sample analysis time of ∼12 min, complete data analysis takes ∼80 h. In contrast, analysis with the GlcR module in the 384-well format is finished within 4 to 8 h, which is at least 10 times faster. In addition, the cost per sample for LC-MS is approximately US$7 (2020 – 2022 average, without instrument purchase included), while IVT-based biosensing reagent costs are approximately US$0.5 for a 20 µL reaction, which makes our system a more cost and resource-efficient alternative to LC-MS measurements (see Supplementary Note 1 for a detailed comparison of methods, and Supplementary Table 7 for IVT reagent costs).

We note that the applicability of our approach depends on the availability of suitable aTF-operator pairs for sensing the metabolite of interest. Today, only a few of allosteric transcription factors are sufficiently characterized and collected in curated databases such as GroovDB^37^. However, systems-biology approaches have proven powerful in identifying the role of aTFs *in vivo*^38–42^, and are complemented recently through computational approaches that were able to successfully predict new aTF-operator pairs^43–46^ and enzymes converting non-detectable metabolites into detectable ones^47,48^. At the same time, efforts to engineer aTFs for new effector specificities and enhanced properties are increasing^42,49–51^, which will hopefully increase the repertoire and availability of aTFs to construct new IVT biosensors in the future.

## Methods

### Chemicals

Unless stated differently, chemicals were purchased from Merck KGaA (Darmstadt, Germany) and Carl Roth GmbH (Karlsruhe, Germany). Commercial enzymes and bioreagents were purchased from New England Biolabs (Frankfurt am Main, Germany).

### Strains and growth media

For molecular cloning, *E. coli* NEB turbo was grown in lysogeny broth supplemented with an appropriate antibiotic (100 µg/mL ampicillin or 34 µg/mL chloramphenicol). For protein production, either *E. coli* M15 (T7 RNA polymerase) or *E. coli* BL21-AI (MBP-GlcR) was grown in terrific broth (TB) supplemented with 100 µg/mL ampicillin or 34 µg/mL chloramphenicol, respectively. All strains used are listed in Supplementary Table 1.

### Assembly of plasmids and preparation of linear DNA templates

Oligonucleotides were purchased from Merck KGaA. Synthetic dsDNA was purchased from Twist Bioscience (South San Francisco, CA, USA). Sanger sequencing was performed by MicroSynth (Göttingen, Germany). Plasmids were generated by Golden Gate Assembly using the Modular Cloning system proposed by Stukenberg et al.^52^ 0.4 nM vector DNA and 4 to 8 nM insert DNA were assembled using 0.5 U/µL Esp3I or 1 U/µL BsaI-HFv2, 40 U/µL T4 ligase in 1x T4 ligase buffer. Reactions were cycled 15 times for 1.5 min at 37°C and 3 min at 16°C. Enzymes were heat-inactivated for 5 min at 50°C and 10 min at 80°C. Golden Gate product was transformed into chemically competent *E. coli* NEB turbo cells, and individual clones were verified by Sanger sequencing using oligonucleotides oSB0021 and oSB022.

All linear DNA templates were prepared by PCR amplification from the respective plasmids using oligonucleotides oSB0021 and oSB0022, and Q5 DNA polymerase, following the vendor’s instructions. All amplified DNA fragments were purified using the NucleoSpin Gel and PCR Clean-up kit (Macherey-Nagel), according to the vendor’s instructions. DNA concentrations were calculated from absorbance measurements at 260 nm (A_260_) using a NanoDrop2000 spectrophotometer (Thermo Scientific, Waltham, MA, USA). To increase the throughput of screening various T7 promoter-operator sequences, we developed a workflow (inspired by previous work^53,54^) which is described in detail in Supplementary Note 2 and the Supplementary Methods. All plasmids, linear templates and oligonucleotides used are listed in Supplementary Table 2-4, respectively.

### Protein production and purification

MBP-GlcR was produced and purified as previously described by Schada von Borzyskowski et al.^29^. T7 RNA polymerase was produced in an *E. coli* M15 strain harboring plasmid pQE30-T7 RNAP^55^ (strain sAP94). First, a pre-culture was inoculated in terrific broth (TB), supplemented with 100 µg/mL ampicillin. Cells were grown to high density overnight at 37°C. The pre-culture was used the following day to inoculate a production culture in TB medium supplemented with 100 µg/mL ampicillin and Antifoam reagent. The culture was grown in a baffled flask at 37°C until an OD_600_ of 0.7 was reached. The culture was then cooled down to room temperature for 30 min, before inducing the culture with 0.5 mM IPTG. Cells were grown overnight at 20°C. Cells were harvested at 4,000 × *g* for 20 min at 12°C, and cell pellets were resuspended in twice their volume of buffer A (50 mM HEPES pH 7.5, 500 mM KCl) with 5 mM MgCl_2_ and DNase I (Roche, Basel, Switzerland). Cells were lysed by sonication using a SonoplusGM200 (BANDELIN electronic GmbH & Co. KG, Berlin, Germany) equipped with a KE76 tip at 50% amplitude for 3x 1 min of 1 s on/off pulses. Lysates were cleared by centrifugation at 100,000 × *g* for 1 h at 8°C, and the supernatant was then filtered through 0.45 µm filters (Sarstedt, Nümbrecht, Germany). For affinity purification, an Äkta Start FPLC system (formerly GE Healthcare, now Cytiva, Marlborough, MA, USA) was used with two stacked 1 mL Ni-NTA columns (HiTrap HP, Cytiva).

The cleared lysate was loaded onto the columns, which were equilibrated with buffer A. The column was washed with buffer A + 75 mM imidazole and eluted with buffer A + 500 mM imidazole. The eluate was desalted using two stacked 5 mL HiTrap desalting columns (Sephadex G-25 resin, Cytiva) and protein elution buffer (25 mM Tris-HCl pH 7.4, 100 mM NaCl). Protein concentration was calculated from absorbance at 280 nm (A_280_) on a NanoDrop2000, and respective extinction coefficients, calculated by ProtParam (https://web.expasy.org/protparam/). Purified T7 RNA polymerase was aliquoted, flash-frozen in liquid nitrogen and stored at -70°C.

### *In vitro* transcription assays

*In vitro* transcription reactions were typically set up, unless stated differently, by adding the following components at their final concentration: IVT buffer (40 mM HEPES (pH 7.8), 8 mM MgCl_2_, 10 mM DTT, 20 mM NaCl and 2 mM spermidine), 0.2 mM DFHBI-1T, 11.4 mM nucleoside triphosphates (rNTPs, Thermo Fisher Scientific), 0.015 U/µL thermostable inorganic pyrophosphatase (New England Biolabs) and 15 nM DNA template. 750 nM MBP-GlcR (MGlcR) were added to the GlcR module. To ensure stability of rNTPs, stocks at 80 mM rNTPs are buffered in 200 mM Tris base. The sample volume of an IVT reaction was 20 µL, prepared in replicates of *n=3*. The reaction mix (57.8 µL = 3.4x 17 µL) was equilibrated at RT for 30 min, before adding the respective analyte in a 1:10 dilution (6.8 µL = 3.4x 2 µL, e.g. effector molecule, individual CETCH components at an indicated concentration or CETCH cycle sample) and 0.33 µM T7 RNA polymerase (3.4 µL = 3.4x 1 µL). The reactions were mixed by pipetting and 3x 20 µL were immediately transferred into a 384-well, black, optically clear, flat-bottom, non-binding microtiter plate (Greiner Bio-One, Kremsmünster, Austria; catalog no.: 781906). Plates were centrifuged for 30 s in a small table-top plate centrifuge (VWR, Radnor, PA, USA) before measurement. Reactions were characterized in triplicates on a plate reader (Infinite M200, Tecan, Männedorf, Switzerland) at 37°C, with 30 s of shaking before each fluorescence read at 472 nm excitation wavelength and 507 nm emission wavelength. Bottom reads of the plate allow for more precise measurements compared to top reads. To convert arbitrary fluorescence measurements to micromolar equivalents of fluorescein (MEF), serial dilutions of a 12.5 µM stock of NIST-traceable fluorescein standard (Invitrogen, catalog no.: F36915) were prepared in dH_2_O and measured alongside each *in vitro* transcription assay. To convert arbitrary fluorescence units in MEF, 1) fluorescein fluorescence (in arbitrary units) was linearly regressed with fluorescein concentrations (in µM), 2) arbitrary fluorescence units were then divided by the slope of the linear fit. See Jung et al.^20^ for a detailed description of MEF standardization.

*In vitro* transcription assays with *E. coli* lysate were prepared from lysate of *E. coli* BL21 Star as previously described^56^. 20 U RNase inhibitor (NEB, #M0314S) were added to the initial titration of lysate in IVT. Lysate samples were MWCO filtered using 3 kDa and 10 kDa Amicon filters (Merck Millipore, catalog no.: UFC500308 (3 kDa), UFC501008 (10 kDa)) for 30 min at 14,000 × *g* and 4°C.

### CETCH cycle assays

The production and purification of enzymes was done as previously described by Sundaram et al.^4^. To test whether stopping reactions by removing enzymes through MWCO filtration yields the same glycolate concentration as stopping reactions by protein precipitation with formic acid, we ran a single CETCH cycle assay (day 7, condition 15^15^) in an 80 µL volume (1.5 mL microcentrifuge tube; started with 100 µM propionyl-CoA substrate; 300 rpm shaking in a thermoshaker for 3 h at 30°C; see concentrations in Supplementary Table 6). Two 9 µL samples were quenched with 1 µL of 50% formic acid and two 25 µL samples were filtered through a 10 kDa MWCO plate (PALL AcroPrep Advance 96-well filter plate; 350 µL, Omega 10K MWCO, catalog no.: 8034) by centrifugation (15 min, 2272 × *g*, 20°C). 2 µL of the samples were diluted in 18 µL of ddH_2_O and used for quantification via LC-MS (method previously described by Pandi et al.^15^).

To generate different glycolate concentrations in constant buffer and cofactor conditions, we ran CETCH cycle assays in which only the methylsuccinyl-CoA oxidase (Mco) concentration was titrated. Six reactions of condition 15 (see Supplementary Table 6) were prepared in a 125 µL volume with different concentrations of Mco: 2x, 1x, 0.5x, 0.25x, 0.1x & no Mco (1x = 26 µM). Reactions were prepared in 1.5 mL microcentrifuge tubes, started with 100 µM propionyl-CoA and shaken for 3 h at 30°C and 300 rpm in a thermoshaker. Samples were filtered and glycolate was quantified as described above. Filtered samples were stored at -20°C.

To prepare CETCH cycle samples with varied buffer and cofactor conditions, samples were prepared in a 150 µL volume (1.5 mL microcentrifuge tube; started with 100 µM propionyl-CoA substrate; 500 rpm shaking in a thermoshaker for 4 h at 30°C). Concentrations of individual CETCH cycle components were varied in the following ranges: HEPES (75 – 200 mM, pH 7.4 – 7.8), MgCl_2_ (2.5 – 17.5 mM), CP (5 – 60 mM), Sodium bicarbonate (2.5 – 60 mM), Sodium formate (10 – 60 mM), CoA (0 – 5 mM), coenzyme B_12_ (0 – 0.1 mM), ATP (3 – 5 mM), NADPH (2.5 – 10 mM), propionyl-CoA oxidase (Pco, 0.10 – 9.57 μM), crotonyl-CoA carboxylase/reductase (Ccr, 0.62 – 2.78 μM), epimerase (Epi, 0.74 – 6.70 μM), methylmalonyl-CoA mutase (Mcm, 0.61 – 2.89 μM), succinyl-CoA reductase (Scr, 3.49 – 13.08 μM), Succinic semialdehyde reductase (Ssr, 0.55 – 4.97 μM), 4-hydroxybutyryl-CoA synthetase (Hbs, 0.53 – 12.28 μM), 4-hydroxybutyryl-CoA dehydratase (Hbd, 0.73 – 3.64 μM), ethylmalonyl-CoA mutase (Ecm, 0.86 – 2.88 μM), methylsuccinyl-CoA oxidase (Mco, 26.01 – 46.54 μM), mesaconyl-CoA hydratase (Mch, 0.28 – 2.84 μM), malyl-CoA/citramalyl-CoA lyase (Mcl1, 2.79 – 14.73 μM), catalase (KatE, 2.46 – 8.21 μM), Formate dehydrogenase (Fdh, 7.28 – 40.77 μM), creatine kinase (CK, 0.78 – 3.14 μM), carbonic anhydrase (CA, 0.02 – 0.13 μM), and glyoxylate/succinic semialdehyde reductase (GOR, 3.31 – 5.25 μM). For a detailed overview of the involved enzymes see Sundaram et al.^4^, and refer to Supplementary Table 6 for details. All assays were started with 0.1 mM propionyl-CoA. After 4 h, samples were filtered through 10 kDa MWCO spin filters (Amicon Ultra 0.5 mL, Merck Millipore, catalog no.: UFC501008), by centrifuging at 14,000 × *g* and 4°C for 20 min. Glycolate from filtrates was quantified as described above (with the minor difference that 10 µM internal ^13^C-glycolate standard was used), and samples were stored at -20°C.

### Overview of the Off-line IVT Biosensing Workflow (all details are described above)

1. prepare and run CETCH samples
2. filter CETCH samples through 10 kDa MWCO membrane (plate or spin column-based)
3. prepare dilution series NIST-traceable fluorescein standard and transfer to 384-well plate
4. prepare ROSALIND reaction mix on ice

a. omit CETCH sample and T7 RNA polymerase
b. prepare in a 3.4x scale to prepare three replicates per sample
5. aliquot 57.8 µL in PCR tube strips, equilibrate at RT for 30 min to ensure good repression by GlcR
6. add 6.8 µL CETCH samples to respective wells → 1:10 dilution of the CETCH sample in the IVT sensor
7. add 3.4 µL T7 RNA polymerase to each well
8. mix by pipetting up and down a volume of 40 µL
9. transfer 20 µL in triplicates in a 384-well plate, centrifuge the plate and start plate reader measurement
10. linearly regress fluorescein standard data to calculate MEF values from arbitrary units for data analysis

## Acknowledgments

The authors would like to thank Lennart Schada von Borzyskowski, Katharina Kremer, Amir Pandi, Blake Rasor and Scott Scholz for helpful discussions; and Peter Claus for technical assistance with the operation of the LC-MS instrument. We also thank Lennart Schada von Borzyskowski and Katharina Kremer for providing the GlcR sequence and intergenic sequence *pden4399-4400*, and Amir Pandi for providing *E. coli* strain sAP94 and his feedback on the manuscript.

S.G. is grateful to the European Molecular Biology Organization (EMBO) postdoctoral fellowship (S.G. ALTF 162-2022). N.B. conducted his research within the Max Planck School Matter to Life supported by the German Federal Ministry of Education and Research (BMBF) in collaboration with the Max Planck Society.

## Author Contributions

Conceptualization, S.B., L.B., and T.J.E.; Methodology, S.B. and L.B.; Investigation, S.B., L.B., C.D., N.B. and S.G.; Visualization: S.B.; Writing – Original Draft, S.B. and T.J.E.; Writing – Review & Editing, S.B. and T.J.E.; Funding Acquisition, T.J.E.; Resources, N.P..; Supervision, S.B. and T.J.E.

## References

1. Bowie, J. U. et al. Synthetic Biochemistry: The Bio-inspired Cell-Free Approach to Commodity Chemical Production. Trends in Biotechnology 38, 766–778 (2020).

2. Schwander, T., Schada von Borzyskowski, L., Burgener, S., Cortina, N. S. & Erb, T. J. A synthetic pathway for the fixation of carbon dioxide in vitro. Science 354, 900 LP – 904 (2016).

3. Luo, S. et al. Construction and modular implementation of the THETA cycle for synthetic CO2 fixation. Nature Catalysis 6, 1228–1240 (2023).

4. Sundaram, S. et al. A Modular In Vitro Platform for the Production of Terpenes and Polyketides from CO2. Angewandte Chemie International Edition 60, 16420–16425 (2021).

5. Diehl, C., Gerlinger, P. D., Paczia, N. & Erb, T. J. Synthetic anaplerotic modules for the direct synthesis of complex molecules from CO2. Nature Chemical Biology 19, 168–175 (2023).

6. Valliere, M. A. et al. A cell-free platform for the prenylation of natural products and application to cannabinoid production. Nature Communications 10, 565 (2019).

7. Valliere, M. A., Korman, T. P., Arbing, M. A. & Bowie, J. U. A bio-inspired cell-free system for cannabinoid production from inexpensive inputs. Nature Chemical Biology 16, 1427–1433 (2020).

8. Korman, T. P., Opgenorth, P. H. & Bowie, J. U. A synthetic biochemistry platform for cell free production of monoterpenes from glucose. Nature Communications 8, 1–8 (2017).

9. Claassens, N. J., Burgener, S., Vögeli, B., Erb, T. J. & Bar-Even, A. A critical comparison of cellular and cell-free bioproduction systems. Current Opinion in Biotechnology 60, 221–229 (2019).

10. Karim, A. S. et al. In vitro prototyping and rapid optimization of biosynthetic enzymes for cell design. Nature Chemical Biology 16, 912–919 (2020).

11. Vögeli, B. et al. Cell-free prototyping enables implementation of optimized reverse β-oxidation pathways in heterotrophic and autotrophic bacteria. Nature Communications 13, 3058 (2022).

12. Liew, F. E. et al. Carbon-negative production of acetone and isopropanol by gas fermentation at industrial pilot scale. Nature Biotechnology (2022) doi:10.1038/s41587-021-01195-w.

13. Dudley, Q. M., Karim, A. S., Nash, C. J. & Jewett, M. C. In vitro prototyping of limonene biosynthesis using cell-free protein synthesis. Metabolic Engineering 61, 251–260 (2020).

14. Kelwick, R. et al. Cell-free prototyping strategies for enhancing the sustainable production of polyhydroxyalkanoates bioplastics. Synthetic Biology 3, (2018).

15. Pandi, A. et al. A versatile active learning workflow for optimization of genetic and metabolic networks. Nature Communications 13, 3876 (2022).

16. McLean, R. et al. Exploring alternative pathways for the in vitro establishment of the HOPAC cycle for synthetic CO2 fixation. Science Advances 9, eadh4299 (2023).

17. Sakai, A. et al. Cell-Free Expression System Derived from a Near-Minimal Synthetic Bacterium. ACS Synthetic Biology (2023) doi:10.1021/acssynbio.3c00114.

18. Morini, L. et al. Leveraging Active Learning to Establish Efficient In Vitro Transcription and Translation from Bacterial Chromosomal DNA. ACS Omega (2024) doi:10.1021/acsomega.4c00111.

19. Miller, T. E. et al. Light-powered CO2 fixation in a chloroplast mimic with natural and synthetic parts. Science 368, 649 LP – 654 (2020).

20. Jung, J. K. et al. Cell-free biosensors for rapid detection of water contaminants. Nature Biotechnology (2020) doi:10.1038/s41587-020-0571-7.

21. Alam, K. K., Tawiah, K. D., Lichte, M. F., Porciani, D. & Burke, D. H. A Fluorescent Split Aptamer for Visualizing RNA–RNA Assembly In Vivo. ACS Synthetic Biology 6, 1710–1721 (2017).

22. Giaveri, S. et al. An interdependent Metabolic and Genetic Network shows emergent properties in vitro. bioRxiv 2023.11.26.568713 (2023) doi:10.1101/2023.11.26.568713.

23. Rothschild, L. J. et al. Building Synthetic Cells─From the Technology Infrastructure to Cellular Entities. ACS Synth. Biol. (2024) doi:10.1021/acssynbio.3c00724.

24. Schwille, P. et al. MaxSynBio: Avenues Towards Creating Cells from the Bottom Up. Angewandte Chemie International Edition 57, 13382–13392 (2018).

25. Pellicer, M. T., Badía, J., Aguilar, J. & Baldomà, L. glc locus of Escherichia coli: characterization of genes encoding the subunits of glycolate oxidase and the glc regulator protein. Journal of Bacteriology 178, 2051–2059 (1996).

26. Pellicer, M. T. et al. Cross-induction of glc and ace Operons of Escherichia coli Attributable to Pathway Intersection: Characterization of the glc promoter. Journal of Biological Chemistry 274, 1745–1752 (1999).

27. Xu, S., Zhang, L., Zhou, S. & Deng, Y. Biosensor-based multi-gene pathway optimization for enhancing the production of glycolate. Applied and Environmental Microbiology AEM.00113-21 (2021) doi:10.1128/AEM.00113-21.

28. Schada von Borzyskowski, L., et al. Marine Proteobacteria metabolize glycolate via the β-hydroxyaspartate cycle. Nature 575, 500–504 (2019).

29. Schada von Borzyskowski, L. et al. Multiple levels of transcriptional regulation control glycolate metabolism in Paracoccus denitrificans. bioRxiv 2024.03.11.584432 (2024) doi:10.1101/2024.03.11.584432.

30. Seo, K. & Ichihashi, N. Investigation of Compatibility between DNA Replication, Transcription, and Translation for in Vitro Central Dogma. ACS Synthetic Biology (2023) doi:10.1021/acssynbio.3c00130.

31. Rasor, B. J. et al. Toward sustainable, cell-free biomanufacturing. Current Opinion in Biotechnology 69, 136–144 (2021).

32. Takahashi, M. K. et al. Rapidly Characterizing the Fast Dynamics of RNA Genetic Circuitry with Cell-Free Transcription–Translation (TX-TL) Systems. ACS Synthetic Biology 4, 503–515 (2015).

33. Boyd, M. A., Thavarajah, W., Lucks, J. B. & Kamat, N. P. Robust and tunable performance of a cell-free biosensor encapsulated in lipid vesicles. Science Advances 9, eadd6605 (2023).

34. Sharon, J. A. et al. Trumpet is an operating system for simple and robust cell-free biocomputing. Nature Communications 14, 2257 (2023).

35. Schoenmakers, L. L. J. et al. In Vitro Transcription–Translation in an Artificial Biomolecular Condensate. ACS Synthetic Biology 12, 2004–2014 (2023).

36. Gonzales, D. T., Yandrapalli, N., Robinson, T., Zechner, C. & Tang, T.-Y. D. Cell-Free Gene Expression Dynamics in Synthetic Cell Populations. ACS Synthetic Biology 11, 205–215 (2022).

37. d’Oelsnitz, S., Love, J. D., Diaz, D. J. & Ellington, A. D. GroovDB: A Database of Ligand-Inducible Transcription Factors. ACS Synthetic Biology 11, 3534–3537 (2022).

38. Lempp, M. et al. Systematic identification of metabolites controlling gene expression in E. coli. Nat Commun 10, 4463 (2019).

39. Donati, S. et al. Multi-omics Analysis of CRISPRi-Knockdowns Identifies Mechanisms that Buffer Decreases of Enzymes in *E. coli* Metabolism. Cell Systems 12, 56–67.e6 (2021).

40. Gagarinova, A. et al. Auxotrophic and prototrophic conditional genetic networks reveal the rewiring of transcription factors in Escherichia coli. Nature Communications 13, 4085 (2022).

41. Rodionova, I. A. et al. A systems approach discovers the role and characteristics of seven LysR type transcription factors in Escherichia coli. Scientific Reports 12, 7274 (2022).

42. Pearson, A. N. et al. Characterization and Diversification of AraC/XylS Family Regulators Guided by Transposon Sequencing. ACS Synthetic Biology 13, 206–219 (2024).

43. Hanko, E. K. R. et al. A genome-wide approach for identification and characterisation of metabolite-inducible systems. Nature Communications 11, 1213 (2020).

44. Hanko, E. K. R., Joosab Noor Mahomed, T. A., Stoney, R. A. & Breitling, R. TFBMiner: A User-Friendly Command Line Tool for the Rapid Mining of Transcription Factor-Based Biosensors. ACS Synthetic Biology 12, 1497–1507 (2023).

45. d’Oelsnitz, S., Ellington, A. D. & Ross, D. J. Ligify: Automated genome mining for ligand-inducible transcription factors. bioRxiv 2024.02.20.581298 (2024) doi:10.1101/2024.02.20.581298.

46. d’Oelsnitz, S., Stofel, S. K., Love, J. D. & Ellington, A. D. Snowprint: a predictive tool for genetic biosensor discovery. Commun Biol 7, 1–9 (2024).

47. Delépine, B., Libis, V., Carbonell, P. & Faulon, J.-L. SensiPath: computer-aided design of sensing-enabling metabolic pathways. Nucleic Acids Research 44, W226–W231 (2016).

48. Pandi, A. et al. Metabolic perceptrons for neural computing in biological systems. Nature Communications 10, 3880 (2019).

49. Snoek, T. et al. Evolution-guided engineering of small-molecule biosensors. Nucleic Acids Research 48, e3–e3 (2020).

50. d’Oelsnitz, S., et al. Using fungible biosensors to evolve improved alkaloid biosyntheses. Nature Chemical Biology (2022) doi:10.1038/s41589-022-01072-w.

51. d’Oelsnitz, S., Nguyen, V., Alper, H. S. & Ellington, A. D. Evolving a Generalist Biosensor for Bicyclic Monoterpenes. ACS Synthetic Biology (2022) doi:10.1021/acssynbio.1c00402.

52. Stukenberg, D. et al. The Marburg Collection: A Golden Gate DNA Assembly Framework for Synthetic Biology Applications in Vibrio natriegens. ACS Synthetic Biology 10, 1904–1919 (2021).

53. Sun, Z. Z., Yeung, E., Hayes, C. A., Noireaux, V. & Murray, R. M. Linear DNA for Rapid Prototyping of Synthetic Biological Circuits in an Escherichia coli Based TX-TL Cell-Free System. ACS Synthetic Biology 3, 387–397 (2014).

54. Lehr, F.-X., et al. Modular Golden Gate Assembly of Linear DNA Templates for Cell-free Prototyping. Preprint at 10.48550/arXiv.2310.13665 (2023).

55. Shimizu, Y. et al. Cell-free translation reconstituted with purified components. Nature Biotechnology 19, 751–755 (2001).

56. Rasor, B. J., Vögeli, B., Jewett, M. C. & Karim, A. S. Cell-Free Protein Synthesis for High-Throughput Biosynthetic Pathway Prototyping. in Methods in Molecular Biology (eds. Karim, A. S. & Jewett, M. C.) 199–215 (Springer US, New York, NY, 2022). doi:10.1007/978-1-0716-1998-8_12.

