## Supplementary Information for "*In vitro* transcription-based biosensing of glycolate for prototyping of a complex enzyme cascade"

DOI:

#### Supplementary Notes

**Supplementary Note 1:** Evaluation of throughput, cost, precision, availability and effort of using IVT-based biosensing versus LC-MS measurement for screening.

In addition to throughput, cost and precision of screening (as mentioned in the main text), other factors need to be considered. These include the availability of instruments and expertise as well as the effort required to establish the respective screening method.

**Throughput:** Our routine quantification of glycolate requires 12 min of measurement time per sample, plus sample preparation for LC-MS measurement: first, quenching with formic acid and centrifugation for 1 h, then two successive dilutions in water and internal standard. In comparison, IVT-based measurements typically take about 4 to 8 h for a plate with up to 384 wells. Sample preparation for ROSALIND assays typically takes 2 h, but can be significantly reduced to about 1 h by automating liquid handling, e.g., using an Echo liquid handler. High-throughput, plate-based MWCO filtration of samples adds another 30 min to the workflow. For example, 384 samples would take approximately 79 h for LC-MS measurement, but only 7 h using IVT-based biosensing, which already includes sample preparation time. In contrast, 20 samples would require 5.5 h for LC-MS and approximately 6 h for IVT-based biosensing. This example highlights the usefulness of IVT-based biosensing for screening, but also shows that it is not applicable for small sample sizes, given the effort required to set up the system.

**Cost:** The cost of measurements must be divided into the cost of reagents and the cost of acquisition, maintenance, and staff. For LC-MS measurements, the reagent cost per measurement is low compared to the acquisition and maintenance costs of LC-MS instruments. Because of the high acquisition cost, LC-MS instruments are often centralized in core facilities that require staff to run the samples. In our core facility, the costs in 2020-2022 were as follows. Full cost (per sample, including consumables, reagents, acquisition, maintenance, and staff): US$7.7 ± 0.77. Excluding staff cost (per sample): US$3.33 ± 0.42. Instrument maintenance and measurement cost only (per sample): US$2.73 ± 0.18. Reagents and consumables cost only (per sample): US$0.9 ± 0.18. Acquisition cost in this period per sample were about US$0.6. Without acquisition cost, the cost per sample is at approximately US$7. This shows that the overhead costs of LC-MS measurements far exceed the actual cost of a measurement. For IVT-based measurements, Jung et al. calculated US$0.67 per sample in 2019 (see SI by Jung et al.[1]). We were able to reduce the cost to approximately US$0.49 per sample by reducing the DFHBI-1T and DNA template concentrations (see Supplementary Table 7), plus approximately US$20 for consumables. Suitable plate readers are widely available in laboratories; new acquisition is assumed to be less than US$50,000; no staff is required. In summary, the reagent and consumable costs for ROSALIND and LC-MS measurements are similar, but the high overhead costs of LC-MS measurements make IVT-based screening a more affordable option for the community.

Note that costs have been converted from EUR to USD using a conversion factor of US$1 = 0.92 EUR (April 10, 2024).

**Precision:** The precision of LC-MS measurements is indisputable high, allowing its wide application for qualitative and quantitative measurement of metabolites from complex samples. In this work, we have shown that IVT-based biosensing can be used for screening complex *in vitro* metabolic systems, but we have also shown matrix effects of CETCH cycle components. Such interferences limit the applicability of the system as they need to be identified and resolved first.

**Availability of instruments and expertise:** As discussed in the cost section, LC-MS instruments are expensive and are therefore often centralized in core facilities to make the best economic use of the instrument, which often includes hiring staff to operate the instruments. Although this increases costs, the expertise available in core facilities allows scientists to measure metabolites without LC-MS expertise. In contrast, the plate readers for IVT measurements are widely available in laboratories and can be operated and maintained without assistance. IVT-based biosensing can be quickly established in the laboratory, especially thanks to the thorough documentation by Jung et al.[1–3].

**Efforts to establish a ROSALIND sensor:** We would like to acknowledge that the main limitation in using IVT-based measurements for screening, is the availability of a suitable allosteric transcription factor for the metabolite of interest. Advances in allosteric transcription factor mining[4,5], database collection[6] and engineering[7–10] are critical to unlocking the full potential of biosensing applications. Following our workflow presented in this work, the generation of an IVT-based sensor takes approximately 2-3 weeks, including purification of the allosteric transcription factor, prototyping of promoter-operator sequences and optimization of sensor composition. Despite progress in reducing this time, the time to establish a ROSALIND sensor exceeds the time to optimize LC-MS protocols.

**Conclusion:** Especially for screening large amounts of samples, IVT-based biosensing has advantages, mainly in throughput and instrument availability, but it also requires time to establish IVT-based sensors and to test for potential matrix effects, e.g. IVT inhibition by individual components.

**Supplementary Note 2:** Description of biosensor prototyping using the ROSALIND system.

We developed a workflow for the time-, labor- and resource-efficient generation of linear DNA templates for the prototyping of promoter-operator sequences and their interaction with a corresponding allosteric transcription factor directly in ROSALIND. The workflow, inspired by previous work[11,12], consists of the following steps. 1) hybridization of oligonucleotides encoding the promoter-operator sequences, 2) cloning of hybridized DNA into a ROSALIND reporter vector using Golden Gate Assembly, 3) direct DNA amplification from the crude Golden Gate Assembly reaction using PCR, 4) DNA purification and quantification, and 5) characterization in ROSALIND. This workflow bypasses the tedious steps of DNA transformation in *E. coli* after Golden Gate Assembly, the growth of liquid cultures, plasmid purification and DNA sequencing of clonal DNA - steps that typically add 3-4 days to the process. Our workflow however allowed us to characterize promoter-operator sequences in 1-2 days, greatly increasing our ability to test different T7 promoter-operator sequences.

In detail, promoter-operator DNA sequences with appropriate overhangs were designed and ordered as oligonucleotides from commercial vendors. Next, the oligonucleotides were hybridized to dsDNA and cloned into a ROSALIND reporter vector using Golden Gate Assembly. We constructed a ROSALIND reporter vector (pTE5417) with an *sfGFP*-dropout sequence[13] for the promoter and ribosome binding site position, the 3WJdB reporter sequence[1] and a T7 terminator. Following the nomenclature of Stukenberg et al.[13], the *sfGFP*-dropout spans parts 2 and 3 of a translational unit – consequently hybridized oligonucleotides encode overhangs for part 2 (forward, GGAG) and part 3 (reverse, CATT). The *sfGFP*-dropout sequence is removed during Golden Gate Assembly and was originally designed as a visual selection marker to distinguish between successful (non-fluorescent) and unsuccessful (GFP-fluorescent) clones after transformation in *E. coli*. We next amplified the linear DNA template from the crude Golden Gate Assembly reaction. We take advantage of the difference in length between the *sfGFP*-dropout amplicon (1489 bp, with oSB0021+22) and the much smaller T7 promoter-operator amplicon (~ 500-550 bp) to amplify only DNA encoding the promoter-operator sequences by adjusting the cycle times to a minimum. As a quality control, the PCR product was checked for correct amplicon size and absence of background amplicon of the *sfGFP*-dropout sequence by agarose gel electrophoresis. The DNA concentration is then calculated from the absorbance at 260 nm ($A_{260}$) using the molecular mass of the linear dsDNA molecule. Typically, a 50 µL PCR reaction is sufficient to generate enough DNA to test the IVT output of a promoter-operator sequence with and without the corresponding allosteric transcription factor and in the presence of an effector molecule at 5 nM DNA concentration in triplicate, given 20 µL ROSALIND reaction volume. See Supplementary Methods for a detailed protocol of the workflow.

#### Supplementary Methods

##### Electrophoretic Mobility Shift Assay (EMSA)

Electrophoretic mobility shift assay samples were prepared by adding the following components, listed at their final concentrations: IVT reaction buffer by Jung et al. (40 mM Tris-HCl, pH 8, 20 mM NaCl, 10 mM DTT, 8 mM $MgCl_2$, 2 mM spermidine)[1], 100 nM DNA fragment, annotated concentrations of MGlcR and glycolate, respectively, and 10% glycerol in a total volume of 10 µL. Samples were incubated for 15 min at RT, before loading on native PAGE analysis on 4-20% pre-cast polyacrylamide gels from Bio-Rad Laboratories (Feldkirchen, Germany; catalog no.: 4561095 (20 µL well) or 4561096 (15 µL well)). Native PAGE was performed with 1x Tris/Glycine buffer (pH 8.3) for 30 min at 200 V. The polyacrylamide gel was stained with 3x GelRed® Nucleic acid stain for 20 min in the dark, washed twice with $dH_2O$ for 5 min and imaged under UV light.
For Figure S1A, the intergenic sequence of pden4399-4400 was amplified from plasmid pTE714_4400/4399_ig using oligonucleotides Pden4400_ig_rv and Pden4399_ig_rv, following the protocol for the preparation of linear DNA templates (see main method section). All other DNA fragments were hybridized by mixing 10 µM of each oligonucleotide in 1x T4 ligase buffer (New England Biolabs), heating the mixture to 95°C, and cooling to ~40°C in a thermoshaker.

##### Variation of Golden Assembly and PCR protocol for prototyping T7 promoter-operator sequences in ROSALIND

As described in Supplementary Notes 2, we established a prototyping workflow for the time and resource-efficient prototyping of T7 promoter-operator sequences. First, oligonucleotides were hybridized as described above for EMSA templates, and diluted to 100 nM in 5 mM Tris-HCl (pH 8.5). Golden Gate Assembly with BsaI-HFv2 was performed, as described in the main method section with 5 nM hybridized oligonucleotides, for at least 15 cycles. Higher concentrations of hybridized oligonucleotides inhibited the downstream PCR. Assembled constructs were amplified by PCR using Q5 DNA polymerase and 0.5 µM oligonucleotides (oSB0021+22) according to the vendor's instructions. For thermocycling, we limited the extension time to a minimum to achieve amplification of the promoter-operator construct only – however, the settings may depend on the thermocycler used. For us, the following protocol worked well: 1 cycle of 98°C for 2 min, 30 cycles of [98°C for 10 s, 66°C for 20 s], 1 cycle of 72°C for 10 s. Amplification was verified by agarose gel electrophoresis. PCR products were then purified and measured as described in the main section. Typically, this workflow yielded 750 ng DNA from a 50 µL PCR reaction.

#### Supplementary Tables

**Supplementary Table 1:** All strains used in this study.

| Strain ID | Background strain | Plasmid & resistance maker# | Use | Reference |
|---|---|---|---|---|
| sAP94 | *Escherichia coli* M15 | pQE30-T7 RNAP, $Amp^R$ | Protein production of T7 RNAP | Shimizu et al., (2001)[14] |
| sMGlcR | *Escherichia coli* BL21-AI | pTE5418, $Cam^R$ | Protein production of MBP-GlcR | Schada von Borzykowski et al., (2024)[15] |

### $Amp^R$, ampicillin resistance; $Cam^R$, chloramphenicol resistance

| Commercial strain | Genotype | Source |
|---|---|---|
| *Escherichia coli* NEB turbo | F' *proA*$^+$*B*$^+$ *lacI*$^q$ *∆lacZM15 /fhuA2 ∆(lac-proAB) glnV galK16 galE15 R(zgb-210::Tn10)*$Tet^S$ *endA1 thi-1 ∆(hsdS-mcrB)5* | *New England Biolabs (Frankfurt am Main, Germany)* |
| *Escherichia coli* BL21-AI | $F^-$ *ompT lon hsdS*$_B$ *(*$r_B^-m_B^-$*) gal dcm araB::T7RNAP-tetA* | Thermo Scientific (Waltham, Massachusetts, USA) |
| *Escherichia coli* M15 | $F^-$ *Φ80ΔlacM15 thi lac*$^-$ *mtl*$^-$ *recA*$^+$ | Qiagen (Hilden, Germany) |

**Supplementary Table 2:** All plasmids used in this study*.

| Plasmid ID | Relevant features# | Reference & Addgene |
|---|---|---|
| pTE5400 | Expression vector for N-terminally His-tagged maltose binding protein (10xHis-MBP) with fusion to a protein of interest, $Cam^R$ | Schada von Borzykowski et al., (2024)[15] |
| pTE5417 | Promoter probe vector with dropout sequence for promoter sequences, 3WJdB reporter, T7 terminator, $Amp^R$ | this study |
| pTE5418 | Expression vector for 10x-MBP-GlcR (MGlcR). GlcR is the gene product of *pden4400* from *Paracoccus denitrificans*, codon optimized for *E. coli*, $Cam^R$ | Schada von Borzykowski et al., (2024)[15] |
| pTE5419 | GlcR-$glcO_{36}$ sensor template: *T7 promoter-*$glcO_{36}$*-3WJdB-T7 terminator*, $Amp^R$ | this study |
| pTE5423 | GlcR-$glcO_{60}$ sensor template: *T7 promoter-*$glcO_{60}$*-3WJdB-T7 terminator*, $Amp^R$ | this study |
| pTE714_4400/4399_ig | Encoding intergenic region between *pden4400* and *pden4399* in *Paracoccus denitrificans*, $Tc^R$ | Schada von Borzykowski et al., (2024)[15] |
| pQE30-T7 RNAP | Expression vector for N-terminally His-tagged T7 RNA polymerase, under T5-*lacO* promoter control, $Amp^R$, $Cam^R$ | Shimizu et al., (2001)[14] [Addgene: #124138] |

### $Amp^R$, ampicillin resistance; $Cam^R$, chloramphenicol resistance; $Tc^R$, tetracycline resistance; dropout sequence: explained in Supplementary Note 2

* Note that GlcR sensor templates encoding $glcO_{32}$ and $glcO_{40-56}$ were only assembled as linear template - no plasmids were prepared. Find explanation in Supplementary Note 2.

**Supplementary Table 3:** All linear dsDNA used in this study.

| Template ID | Description | PCR template | Oligonucleotides | Used in |
|---|---|---|---|---|
| ig_*pden4399/4400* | intergenic sequence between *pden4399/4400* | pTE714_4400/4399_ig | Pden4400_ig_rv, Pden4399_ig_rv | Supp. Figure 1A |
| ig_F1 | Fragment 1 of ig_*pden4399/4400* | hybridized | oLUB011+12 | Supp. Figure 1B |
| ig_F2 | Fragment 2 of ig_*pden4399/4400* | hybridized | oLUB013+14 | Supp. Figure 1B |
| ig_F3 | Fragment 3 of ig_*pden4399/4400* | hybridized | oLUB015+16 | Supp. Figure 1B |
| ig_F4 | Fragment 4 of ig_*pden4399/4400* | hybridized | oLUB017+18 | Supp. Figure 1B |
| ig_F5 | Fragment 5 of ig_*pden4399/4400* | hybridized | oLUB019+20 | Supp. Figure 1B |
| ig_F6 | Fragment 6 of ig_*pden4399/4400* | hybridized | oLUB021+22 | Supp. Figure 1B |
| ig_F3.1_20 | Fragment 3.1, 20 bp | hybridized | oLUB029+30 | Supp. Figure 1C |
| ig_F3.2_20 | Fragment 3.2, 20 bp | hybridized | oLUB031+32 | Supp. Figure 1C |
| ig_F3.3_20 | Fragment 3.3, 20 bp | hybridized | oLUB033+34 | Supp. Figure 1C |
| ig_F3.4_20 | Fragment 3.4, 20 bp | hybridized | oLUB035+36 | Supp. Figure 1C |
| ig_F3.5_20 | Fragment 3.5, 20 bp | hybridized | oLUB037+38 | Supp. Figure 1C |
| ig_F3.1_30 | Fragment 3.1, 30 bp | hybridized | oLUB039+40 | Supp. Figure 1C |
| ig_F3.2_30 | Fragment 3.2, 30 bp | hybridized | oLUB041+42 | Supp. Figure 1C |
| ig_F3.3_30 | Fragment 3.3, 30 bp | hybridized | oLUB043+44 | Supp. Figure 1C |
| $P_{T7}$-$glcO_{60}$ | T7 promoter with $glcO_{60}$ | hybridized | oLUB045+46 | Supp. Figure 1D |
| $P_{T7}$-$glcO_{56}$ | T7 promoter with $glcO_{56}$ | hybridized | oLUB047+48 | Supp. Figure 1D |
| $P_{T7}$-$glcO_{52}$ | T7 promoter with $glcO_{52}$ | hybridized | oLUB049+50 | Supp. Figure 1D |
| $P_{T7}$-$glcO_{48}$ | T7 promoter with $glcO_{48}$ | hybridized | oLUB051+52 | Supp. Figure 1D |
| $P_{T7}$-$glcO_{44}$ | T7 promoter with $glcO_{44}$ | hybridized | oLUB053+54 | Supp. Figure 1D |
| $P_{T7}$-$glcO_{40}$ | T7 promoter with $glcO_{40}$ | hybridized | oLUB055+56 | Supp. Figure 1D |
| $P_{T7}$-$glcO_{36}$ | T7 promoter with $glcO_{36}$ | hybridized | oLUB057+58 | Supp. Figure 1D |
| $P_{T7}$-$glcO_{32}$ | T7 promoter with $glcO_{32}$ | hybridized | oLUB059+60 | Supp. Figure 1D |
| $P_{T7}$-$glcO_{60}$ | GlcR sensor with $glcO_{60}$ | prototyping workflow# | oLUB045+46 | Supp. Figure 2A |
| $P_{T7}$-$glcO_{56}$-3WJdB | GlcR sensor with $glcO_{56}$ | prototyping workflow# | oLUB047+48 | Supp. Figure 2A |
| $P_{T7}$-$glcO_{52}$-3WJdB | GlcR sensor with $glcO_{52}$ | prototyping workflow# | oLUB049+50 | Supp. Figure 2A |
| $P_{T7}$-$glcO_{48}$-3WJdB | GlcR sensor with $glcO_{48}$ | prototyping workflow# | oLUB051+52 | Supp. Figure 2A |
| $P_{T7}$-$glcO_{44}$-3WJdB | GlcR sensor with $glcO_{44}$ | prototyping workflow# | oLUB053+54 | Supp. Figure 2A |
| $P_{T7}$-$glcO_{40}$-3WJdB | GlcR sensor with $glcO_{40}$ | prototyping workflow# | oLUB055+56 | Supp. Figure 2A |
| $P_{T7}$-$glcO_{36}$-3WJdB | GlcR sensor with $glcO_{36}$ | prototyping workflow# | oLUB057+58 | Supp. Figure 2A |
| $P_{T7}$-$glcO_{32}$-3WJdB | GlcR sensor with $glcO_{32}$ | prototyping workflow# | oLUB059+60 | Supp. Figure 2A |
| $P_{T7}$-$glcO_{60}$-3WJdB | GlcR sensor with $glcO_{60}$ | pTE5423 | oSB0021+22 | Supp. Figure 2B,C |
| **GlcR sensor template ($P_{T7}$-$glcO_{36}$-3WJdB)** | GlcR sensor with $glcO_{36}$ | pTE5419 | oSB0021+22 | Figure 1-4, Supp. Figure 2B-D, 3-11 |

### see details about prototyping workflow in Supplementary Note 2 and Supplementary Methods

**Supplementary Table 4:** All oligonucleotides used in this study.

| Oligo ID | Sequence [5' to 3'][#] |
|---|---|
| oLUB011 | GGGCGCTCATGCGGTTGTCGGATCCCTGCCATGTAAGCACAGGACGGGCGTGTCG |
| oLUB012 | CGACACGCCCGTCCTGTGCTTACATGGCAGGGATCCGACAACCGCATGAGCGCCC |
| oLUB013 | CTGCCATGTAAGCACAGGACGGGCGTGTCGGTCCAGAAAAATACCGATTGACTAATGTGG |
| oLUB014 | CCACATTAGTCAATCGGTATTTTTCTGGACCGACACGCCCGTCCTGTGCTTACATGGCAG |
| oLUB015 | GTCCAGAAAAATACCGATTGACTAATGTGGTCTGAAAATTATACCAAGAATGCAGGGCAG |
| oLUB016 | CTGCCCTGCATTCTTGGTATAATTTTCAGACCACATTAGTCAATCGGTATTTTTCTGGAC |
| oLUB017 | TCTGAAAATTATACCAAGAATGCAGGGCAGGACACGGGCCTGCCGTTCGACCAAGGGAGG |
| oLUB018 | CCTCCCTTGGTCGAACGGCAGGCCCGTGTCCTGCCCTGCATTCTTGGTATAATTTTCAGA |
| oLUB019 | GACACGGGCCTGCCGTTCGACCAAGGGAGGAGAGCTTGTCCGGTATTGCCATGCCCCGGC |
| oLUB020 | GCCGGGGCATGGCAATACCGGACAAGCTCTCCTCCCTTGGTCGAACGGCAGGCCCGTGTC |
| oLUB021 | AGAGCTTGTCCGGTATTGCCATGCCCCGGCCCGATGCGGGCATCCTGGCGCGTGCGTGA |
| oLUB022 | TCACGCACGCGCCAGGATGCCCGCATCGGGCCGGGGCATGGCAATACCGGACAAGCTCT |
| oLUB029 | GTCCAGAAAAATACCGATTG |
| oLUB030 | CAATCGGTATTTTTCTGGAC |
| oLUB031 | ATACCGATTGACTAATGTGG |
| oLUB032 | CCACATTAGTCAATCGGTAT |
| oLUB033 | ACTAATGTGGTCTGAAAATT |
| oLUB034 | AATTTTCAGACCACATTAGT |
| oLUB035 | TCTGAAAATTATACCAAGAA |
| oLUB036 | TTCTTGGTATAATTTTCAGA |
| oLUB037 | ATACCAAGAATGCAGGGCAG |
| oLUB038 | CTGCCCTGCATTCTTGGTAT |
| oLUB039 | GTCCAGAAAAATACCGATTGACTAATGTGG |
| oLUB040 | CCACATTAGTCAATCGGTATTTTTCTGGAC |
| oLUB041 | GATTGACTAATGTGGTCTGAAAATTATACC |
| oLUB042 | GGTATAATTTTCAGACCACATTAGTCAATC |
| oLUB043 | TCTGAAAATTATACCAAGAATGCAGGGCAG |
| oLUB044 | CTGCCCTGCATTCTTGGTATAATTTTCAGA |
| oLUB045 | ggagTAATACGACTCACTATAGGGGTCCAGAAAAATACCGATTGACTAATGTGGTCTGAAAATTATACCAAGAATGCAGGGCAG |
| oLUB046 | cattCTGCCCTGCATTCTTGGTATAATTTTCAGACCACATTAGTCAATCGGTATTTTTCTGGACCCCTATAGTGAGTCGTATTA |
| oLUB047 | ggagTAATACGACTCACTATAGGGAGAAAAATACCGATTGACTAATGTGGTCTGAAAATTATACCAAGAATGCAGGGCAG |
| oLUB048 | cattCTGCCCTGCATTCTTGGTATAATTTTCAGACCACATTAGTCAATCGGTATTTTTCTCCCTATAGTGAGTCGTATTA |
| oLUB049 | ggagTAATACGACTCACTATAGGGAAATACCGATTGACTAATGTGGTCTGAAAATTATACCAAGAATGCAGGGCAG |
| oLUB050 | cattCTGCCCTGCATTCTTGGTATAATTTTCAGACCACATTAGTCAATCGGTATTTCCCTATAGTGAGTCGTATTA |
| oLUB051 | ggagTAATACGACTCACTATAGGGACCGATTGACTAATGTGGTCTGAAAATTATACCAAGAATGCAGGGCAG |
| oLUB052 | cattCTGCCCTGCATTCTTGGTATAATTTTCAGACCACATTAGTCAATCGGTCCCTATAGTGAGTCGTATTA |
| oLUB053 | ggagTAATACGACTCACTATAGGGATTGACTAATGTGGTCTGAAAATTATACCAAGAATGCAGGGCAG |
| oLUB054 | cattCTGCCCTGCATTCTTGGTATAATTTTCAGACCACATTAGTCAATCCCTATAGTGAGTCGTATTA |
| oLUB055 | ggagTAATACGACTCACTATAGGGACTAATGTGGTCTGAAAATTATACCAAGAATGCAGGGCAG |
| oLUB056 | cattCTGCCCTGCATTCTTGGTATAATTTTCAGACCACATTAGTCCCTATAGTGAGTCGTATTA |
| oLUB057 | ggagTAATACGACTCACTATAGGGATGTGGTCTGAAAATTATACCAAGAATGCAGGGCAG |
| oLUB058 | cattCTGCCCTGCATTCTTGGTATAATTTTCAGACCACATCCCTATAGTGAGTCGTATTA |
| oLUB059 | ggagTAATACGACTCACTATAGGGGGTCTGAAAATTATACCAAGAATGCAGGGCAG |
| oLUB060 | cattCTGCCCTGCATTCTTGGTATAATTTTCAGACCCCCTATAGTGAGTCGTATTA |
| oSB0021 | CGGTTCCTGGCCTTTTGC |
| oSB0022 | GATAGGTGCCTCACTGATTAAGC |
| Pden4399_ig_rv | GATAT**GAATTC**TCACTGGGCCACGGCCTCG |
| Pden4400_ig_rv | CTT**TCTAGA**TCACGCACGCGCCAGGATGCC |

[#] Nucleotides in bold and underlined are recognition sites for endonuclease restriction enzymes, used in Schada von Borzykowski et al.[15] , small letters indicate overhangs after hybridization for downstream Golden Gate Assembly.

**Supplementary Table 5:** Tested concentrations of CETCH cycle components for inhibition of constitutive IVT reactions (see Figure 2 and Supplementary Figure 7).

| CETCH cycle component | Tested concentrations |
|---|---|
| ATP | 0, 0.5, 1, 2, 4, 6, 8, 10 mM |
| B12 | 0, 10, 20, 40, 60, 80, 100, 200, 400 µM |
| 2-mercapoethanol (β-ME) | 0, 1, 2, 4 mM |
| Bicarbonate | 0, 2.5, 5, 10, 20, 40, 60, 80 mM |
| Coenzyme A (CoA) | 0, 0.1, 0.2, 0.3, 0.4, 0.5, 1, 2, 3, 4, 5 mM |
| Creatine phosphate (CP) | 0, 2.5, 5, 10, 20, 40, 60 mM |
| Flavin adenine dinucleotide (FAD) | 0, 10, 20, 40, 60, 80, 100 µM |
| Formate | 0, 2.5, 5, 10, 20, 40, 60, 80 mM |
| Glycerol | 0, 10, 20, 30% (v/v) |
| HEPES | 0, 50, 100, 150, 200 mM |
| $MgCl_2$ | 0, 2.5, 5, 7.5, 10, 12.5, 15, 17.5, 20 mM |
| NADPH | 0, 1, 2, 4, 6, 8, 10 mM |
| Propionyl-CoA | 0, 50, 100, 500 µM |

**Supplementary Table 6:** CETCH cycle conditions used in Figure 3 and 4.

| CETCH component | Sample #1 Day 7 cond. 15# | Sample #2 Day 7 cond. 8# | Sample #3 Day 7 cond. 42# | Sample #4 Day 7 cond. 60# | Sample #5 Day 7 cond. 104# | Sample #6 Day 7 cond. 71# |
|---|---|---|---|---|---|---|
| μM Pco | 3.06 | 9.57 | 1.72 | 3.06 | 5.36 | 0.10 |
| μM Ccr | 1.85 | 1.54 | 2.16 | 0.62 | 2.47 | 2.78 |
| μM Epi | 0.74 | 0.74 | 5.96 | 6.70 | 5.21 | 3.72 |
| μM Mcm | 2.89 | 0.61 | 1.22 | 2.13 | 2.89 | 1.52 |
| μM Scr | 3.49 | 5.23 | 5.23 | 13.08 | 6.97 | 3.49 |
| μM Ssr | 1.66 | 2.21 | 0.55 | 2.21 | 1.66 | 4.97 |
| μM Hbs | 0.53 | 1.60 | 1.60 | 1.60 | 0.53 | 12.28 |
| μM Hbd | 0.73 | 2.18 | 2.91 | 1.45 | 0.73 | 3.64 |
| μM Ecm | 1.44 | 0.86 | 1.73 | 1.15 | 2.59 | 2.88 |
| μM Mco | 26.01 | 46.54 | 26.01 | 26.01 | 26.01 | 34.22 |
| μM Mch | 0.28 | 2.27 | 1.99 | 0.85 | 2.84 | 0.28 |
| μM Mcl1 | 3.58 | 2.79 | 14.73 | 6.37 | 2.79 | 14.73 |
| μM KatE | 3.28 | 6.57 | 8.21 | 3.28 | 2.46 | 6.57 |
| μM Fdh | 30.58 | 30.58 | 23.30 | 40.77 | 7.28 | 7.28 |
| μM CA | 0.07 | 0.02 | 0.02 | 0.08 | 0.13 | 0.03 |
| μM Gor | 4.97 | 5.25 | 3.31 | 4.97 | 3.59 | 4.69 |
| μM CK | 0.78 | 2.35 | 2.75 | 1.96 | 3.14 | 1.18 |
| mM HEPES* | 75 (pH 7.8) | 125 (pH 7.8) | 100 (pH 7.4) | 125 (pH 7.8) | 125 (pH 7.6) | 200 (pH 7.8) |
| mM $MgCl_2$ | 12.5 | 2.5 | 17.5 | 5 | 12.5 | 5 |
| mM CP | 60 | 5 | 5 | 10 | 20 | 20 |
| mM Na bicarbonate | 2.5 | 20 | 60 | 5 | 5 | 5 |
| mM Na formate | 20 | 60 | 20 | 60 | 10 | 60 |
| mM CoA | 0.4 | 0.5 | 0.2 | 0.2 | 5 | 0 |
| mM $B_{12}$ | 0 | 0 | 0 | 0 | 0 | 0.1 |
| mM ATP | 3 | 3 | 3 | 3 | 3 | 5 |
| mM NADPH | 3.75 | 7.5 | 3.75 | 7.5 | 2.5 | 10 |
| mM propionyl-CoA | 0.1 | 0.1 | 0.1 | 0.1 | 0.1 | 0.1 |

* HEPES buffer was pH adjusted by titration with KOH
### condition day + numbers refer to the previously tested conditions by Pandi et al., 2022[16]

**Supplementary Table 7: Cost calculation of a ROSALIND reaction**

| Component | Supplier | Catalog No. | Quantity | Price (USD) | Cost USD/20 µL reaction |
|---|---|---|---|---|---|
| 40 mM HEPES | Carl Roth | 6763.2 | 500 g | $ 141.55 | << $ 0.01 |
| 8 mM $MgCl_2$ | Carl Roth | HN03.2 | 500 g | $ 30.88 | << $ 0.01 |
| 10 mM DTT | Carl Roth | 6908.2 | 25 g | $ 39.57 | << $ 0.01 |
| 20 mM NaCl | Carl Roth | 962.1 | 1000 g | $ 10.47 | << $ 0.01 |
| 2 mM Spermidine | TCI | S2626-1G | 1 g | $ 33.52 | < $ 0.01 |
| 2.85 mM NTP (each) | Thermo Scientific | R1481 | 0.025 mmol each | $ 117.39 | $ 0.25 |
| 0.3 U TIPP | NEB | M0296S | 250 units | $ 68.67 | $ 0.08 |
| 0.2 mM DFHBI-1T | Sigma | SML2697-25MG | 0.025 g | $ 439.36 | $ 0.02 |
| T7 RNAP | Homemade[#] | | ~ 30 nmol (5000 reactions) | $ ~100 | $ 0.02 |
| MGlcR | Homemade[#] | | ~300 nmol (>20,000 reactions) | $ ~ 80 | < $ 0.01 |
| 15 nM DNA template | Homemade[&] | | 50 pmol | $ 18.36 | $ 0.11 |
| **Total** | | | | | **~$ 0.49** |

[#] Homemade protein purification: estimated costs of consumables for purification of the respective protein using the protocol described in the methods.
[&] Homemade linear PCR template: Calculated for the cost of a 0.4 mL PCR and purification using NEB #M0492S ($11.13/prep), oligonucleotides ($0.20/prep) and Macherey-Nagel # 740609.240C ($7.01/prep).
* Prices were calculated using MPI institutional pricing (January 2024), thereby not including shipping, labor, instruments and equipment, consumables and general overhead (electricity, storage).

#### Supplementary Figures

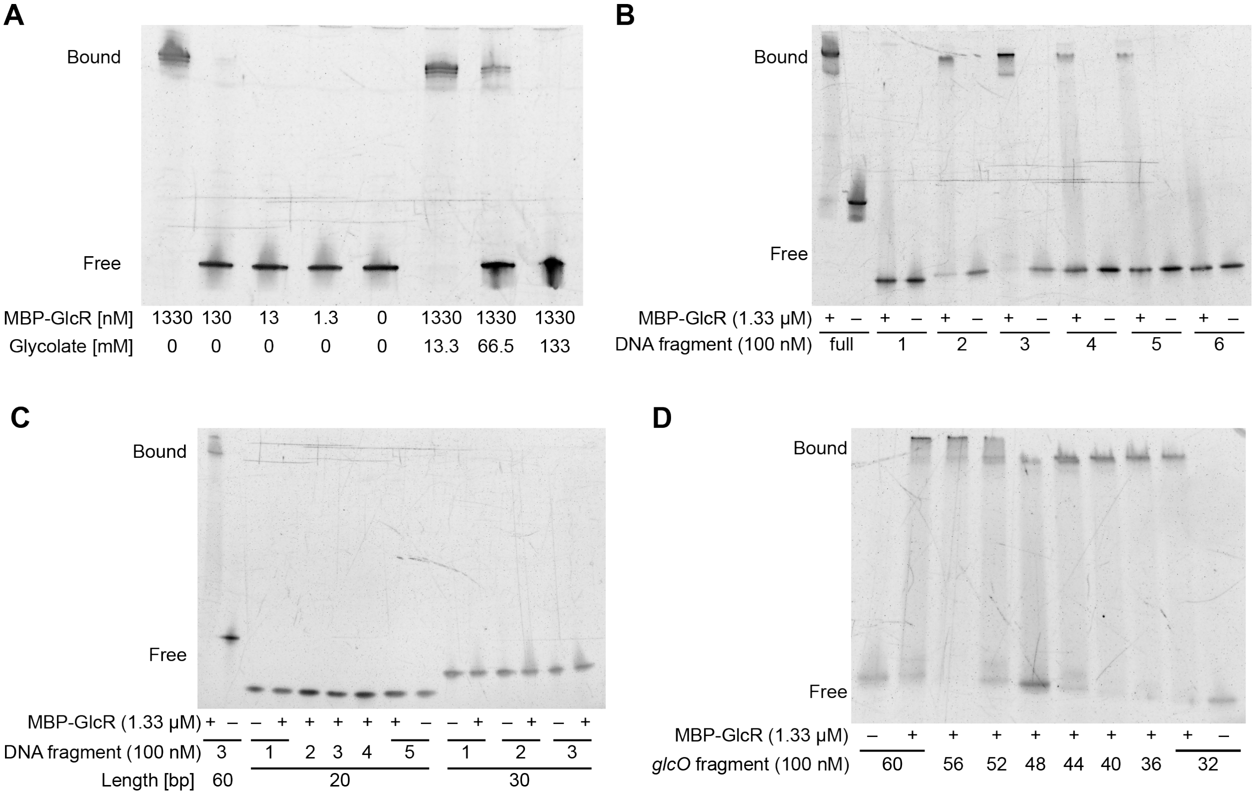


**Supplementary Figure 1:** Electrophoretic mobility shift assays (EMSAs) to investigate the binding of MGlcR to the complete or fragmented *pden4399-4400* intergenic sequence. **A:** Verification of MGlcR binding to the intergenic sequence by titration of MGlcR in the absence of glycolate, and MGlcR unbinding of the intergenic sequence by titration of glycolate with 1330 nM MGlcR. The intergenic sequence concentration is constant at 80.6 nM in each sample. **B:** Search for the operator site of GlcR in the intergenic sequence. Binding of MGlcR to six fragments (55-60 bp, 30 bp overlap) of the intergenic sequence shows binding to fragments #2, #3, #4 and #5. Fragment #3 is bound most strongly. **C**: Narrowing down of the operator site in fragment #3 (see panel B). Binding of MGlcR to eight fragments (5x 20 bp, 3x 30 bp) but no binding is observed. **D:** Binding of MGlcR to 5' truncated sequences of fragment #3. The sequences were truncated in 4 bp steps and suggest binding sites on both flanks of fragment #3. The same fragments were also tested for repression in the ROSALIND system and showed similar results (Supplementary Figure 2A).

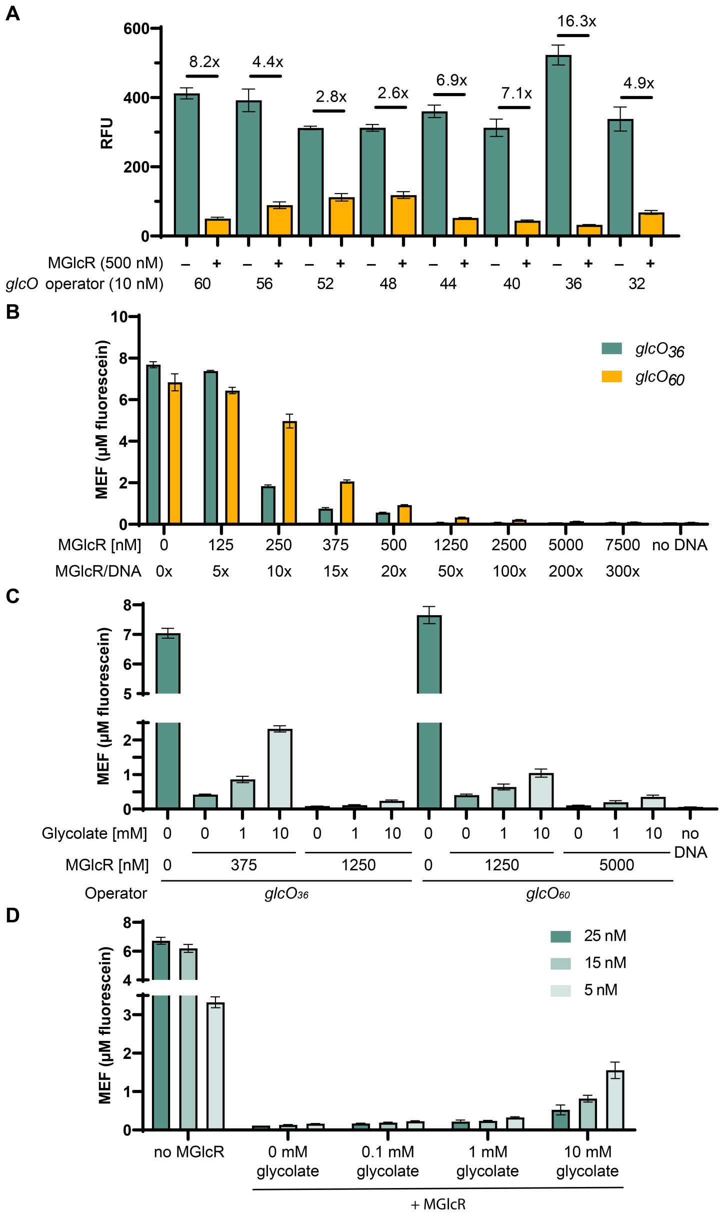


**Supplementary Figure 2:** Construction and optimization of the GlcR sensor module. **A**: Eight $P_{T7}$-*glcO-3WJdB* constructs at 10 nM were assayed for constitutive expression (green) and repression in the presence of 500 nM MGlcR (orange). Operator sequences *glcO*$_{36}$ and *glcO*$_{60}$ showed the most promising results. **B**: $P_{T7}$-*glcO-3WJdB* constructs encoding *glcO*$_{36}$ (green) and *glcO*$_{60}$ (orange) were assayed for the MGlcR/DNA ratio with optimal repression. **C**: Response of MGlcR sensors to 1 and 10 mM glycolate. Sensors are based on *glcO*$_{36}$ with 375 and 1250 nM MGlcR and on *glcO*$_{60}$ with 1250 and 5000 nM MGlcR. **D**: DNA template titration of the *glcO*$_{36}$ sensor at a constant MGlcR/DNA ratio of 50x, namely 25 nM, 15 nM and 5 nM DNA template with 1250 nM, 750 nM and 250 nM MGlcR, respectively, to increase the sensitivity of the sensors to glycolate. Raw fluorescence data are standardized to MEF (µM fluorescein). All data are time points at 4 hours, and are presented as the mean of *n*=3 technical replicates ± s.d. Each IVT reaction was templated with 25 nM linear DNA unless indicated differently.

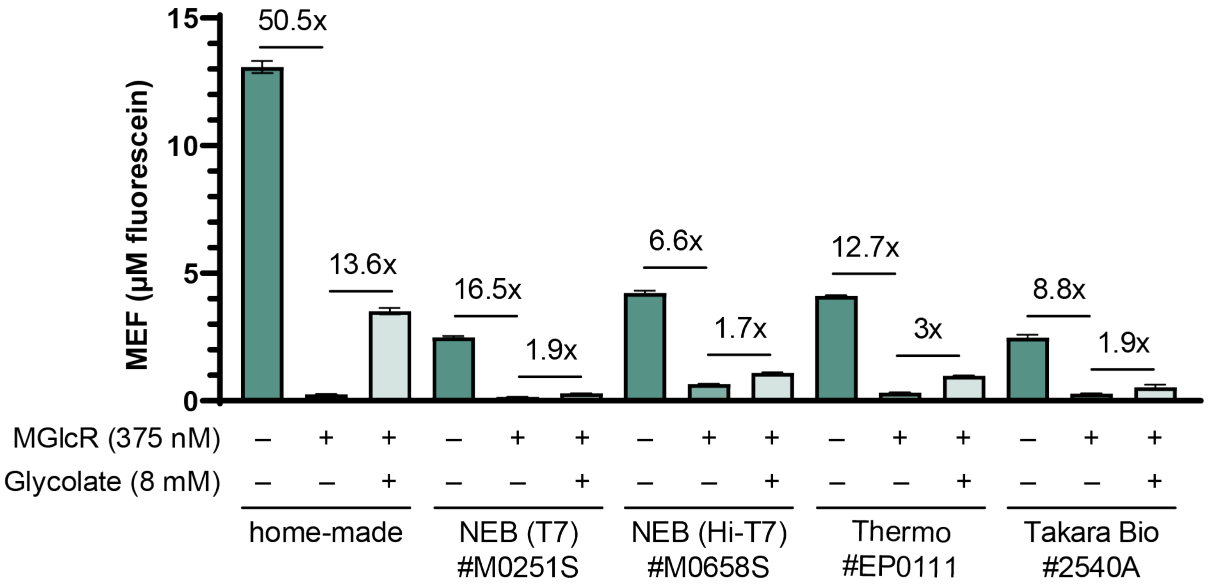


**Supplementary Figure 3:** Testing four commercial T7 RNA polymerases from NEB (catalog no. M0251S & M0658S, 50 units), Thermo Scientific (catalog no. EP0111, 20 units) and Takara Bio (catalog no. 2540A, 50 units) on 1) constitutive expression, 2) repression by MGlcR and 3) de-repression by glycolate. All commercial enzymes show ~3x lower output than homemade T7 RNA polymerase (9.1 pmol). Note that T7 RNA polymerase from Thermo Scientific shows similar constitutive output compared to the other vendors' enzymes despite lower indicated unit concentration. Hi-T7 from NEB shows worse repression by MGlcR in our conditions than all other polymerases. Raw fluorescence data are standardized to MEF (µM fluorescein). Data are the mean of *n*=3 technical replicates ± s.d. of 4 h time points. All IVT reactions were performed in 40 mM HEPES buffer pH 7.8 and were templated by 15 nM linear GlcR sensor template DNA.

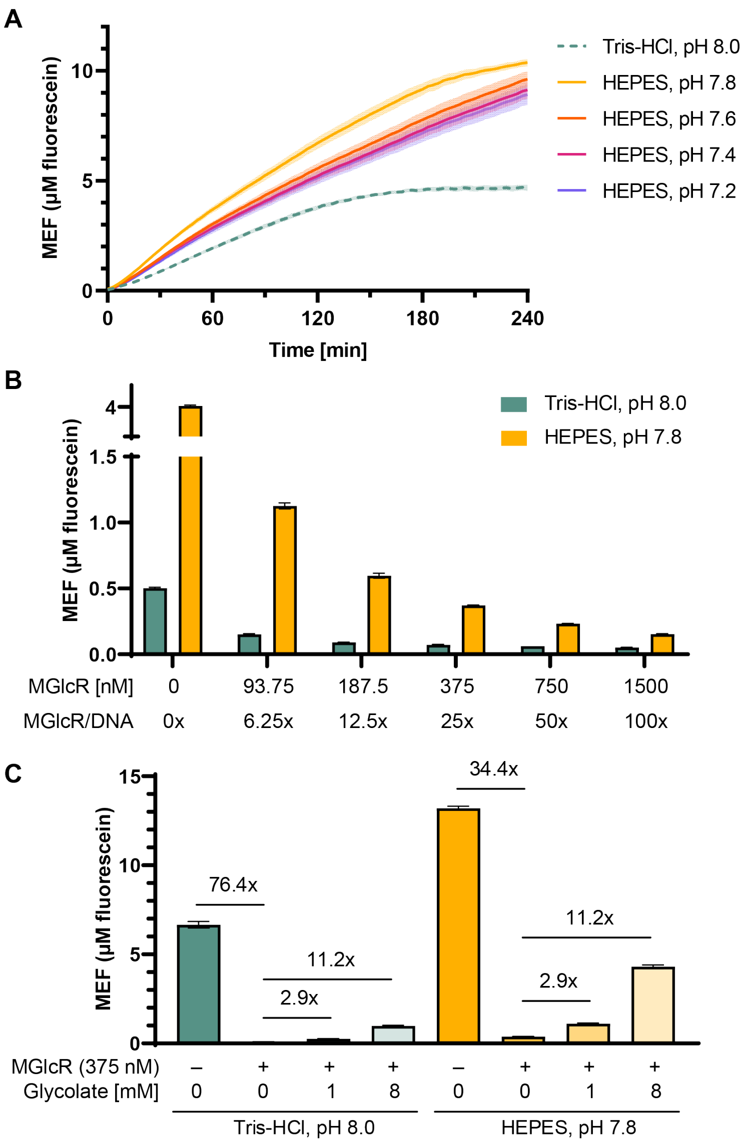


**Supplementary Figure 4:** Comparison of HEPES and Tris-HCl buffers in the ROSALIND system. **A:** pH titration of HEPES buffer from 7.2 to 7.8 and comparison with Tris-HCl pH 8.0 in constitutive IVT. All four HEPES buffer conditions show improved IVT output over Tris-HCl buffer pH 8.0. HEPES buffer pH 7.8 shows the best output. **B:** Titration of MGlcR over DNA template in HEPES buffer pH 7.8 and Tris-HCl buffer pH 8.0. The overall increase in IVT output in HEPES buffer pH 7.8 is accompanied by an increase in background signal. This experiment was carried out with 100 units of T7 RNA polymerase from New England Biolabs (NEB) per reaction. **C:** The glycolate sensor shows similar repression by 375 nM MGlcR and de-repression in the presence of 1 mM or 8 mM glycolate in Tris-HCl pH 8.0 and HEPES pH 7.8, despite the generally increased output in HEPES buffer pH 7.8. Raw fluorescence data are standardized to MEF (µM fluorescein). Data are the mean of *n*=3 technical replicates ± s.d. of a 4 h time course (**A**) or 4 h time points (**B,C**). All IVT reactions were performed in 40 mM buffer and were templated with 15 nM linear GlcR sensor DNA.

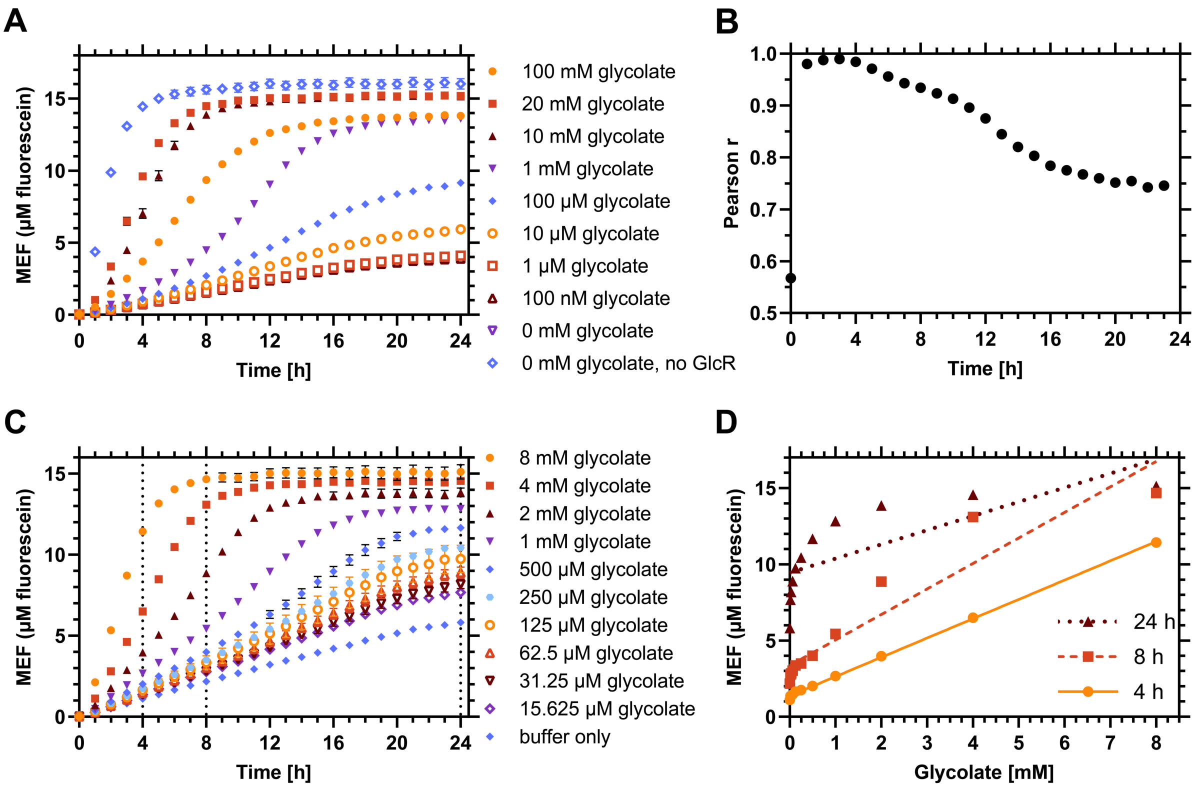


**Supplementary Figure 5:** Dose response of the GlcR sensor module to glycolate. **A:** Titration of glycolate from 100 nM to 100 mM and measurement over 24 hours. Dose-dependent response to up to 20 mM glycolate. Increasing the glycolate concentration to 100 mM inhibits the system. **B:** Pearson correlation coefficients of dose response from 100 nM to 20 mM glycolate were best for the first 4 h and dropped afterwards. **C, D:** Titration of glycolate from 15.625 µM to 8 mM showed a linear dose response after 4 hours when none of the reactions plateaued. As soon as the first reactions plateau, the linearity of the dose response is compromised as shown for 8 and 24 h time points. Raw fluorescence data are standardized to MEF (µM fluorescein). Data are the mean of *n*=3 technical replicates ± s.d. of a 24 h time course (**A, C**) or respectively annotated time ppoints (**B, D**). All IVT reactions were performed in 40 mM HEPES buffer pH 7.8 and templated with 15 nM linear GlcR sensor DNA.

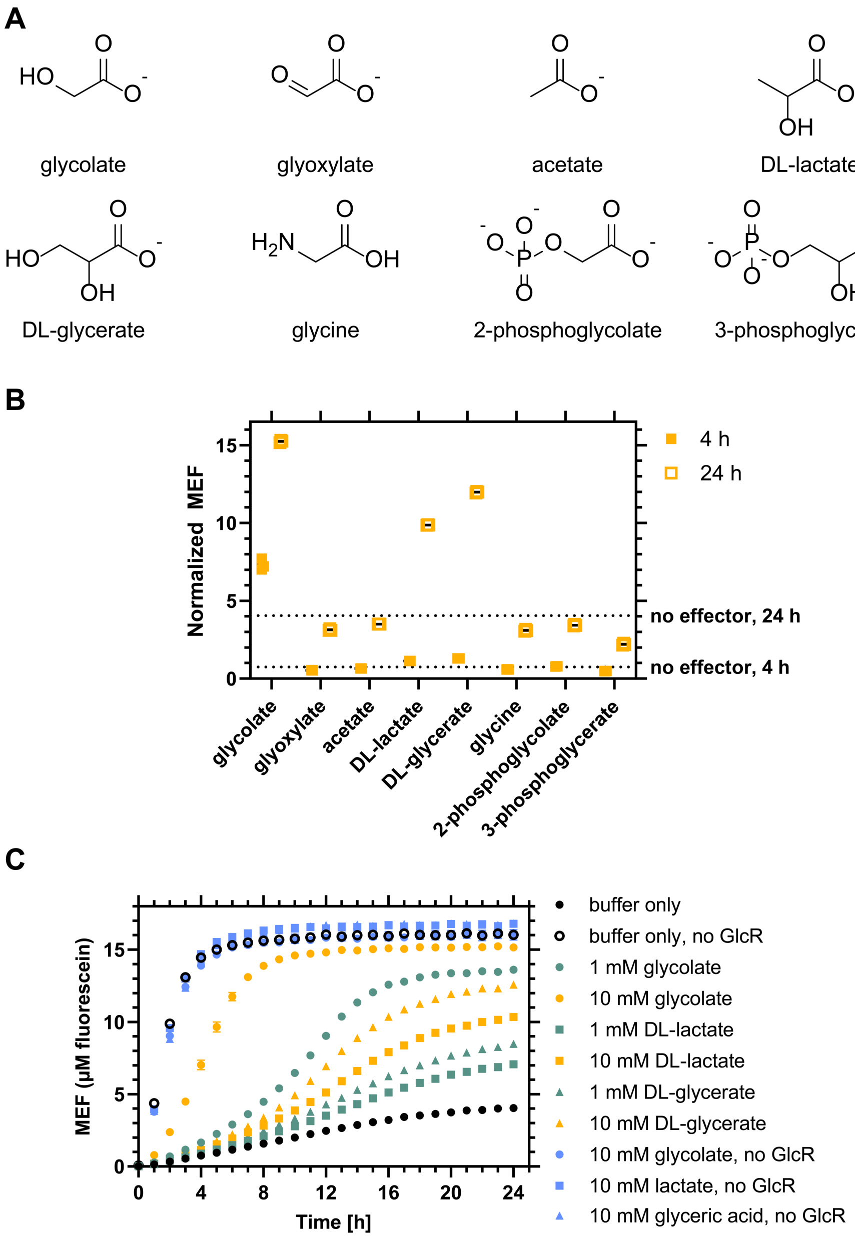


**Supplementary Figure 6:** Promiscuity of GlcR towards structure and context-related molecules. **A:** Chemical structures of assayed molecules. **B:** Response of the GlcR sensor towards 10 mM potential effector molecules after 4 and 24 h. Values are corrected by the effect of the respective molecule in a constitutive IVT reaction, as shown in Figure 1D (blue). See Methods for correction procedure. Lactate and glycerate show a promiscuous effect. **C:** Time course over 24 hours with glycolate, glycerate and lactate at 1 mM (green) and 10 mM (orange), buffer only (black) and in the presence of 10 mM respective effector without MGlcR (blue). Raw fluorescence data are standardized to MEF (µM fluorescein). Data are the mean of *n*=3 technical replicates ± s.d. of a 24 h time course. All IVT reactions were performed in 40 mM HEPES buffer pH 7.8 and templated with 15 nM linear GlcR sensor DNA.

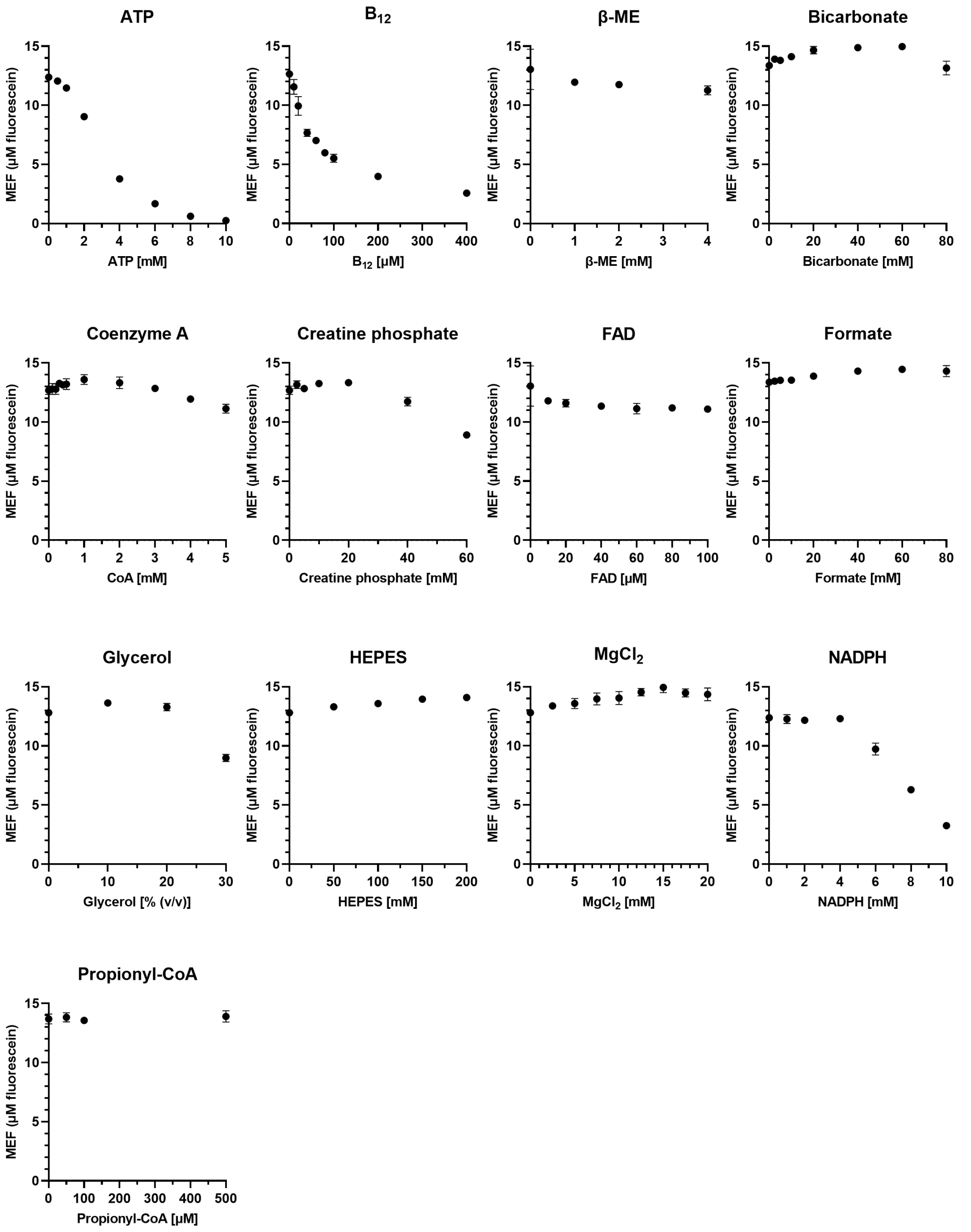


**Supplementary Figure 7:** Titration of CETCH cycle components in constitutive IVT reactions (in absence of GlcR) to investigate their effect on the IVT output. All components were diluted in 50 mM HEPES pH 7.8. Raw fluorescence data are standardized to MEF (µM fluorescein). Data are the mean of *n*=3 technical replicates ± s.d. of 4 h time points. All IVT reactions were performed in 40 mM HEPES buffer (pH 7.8) and were templated with 15 nM linear GlcR sensor DNA. The same data for ATP, B12, NADPH and $MgCl_2$ are shown in Figure 2B. All data are shown normalized to a condition without CETCH cycle component in Figure 2C.

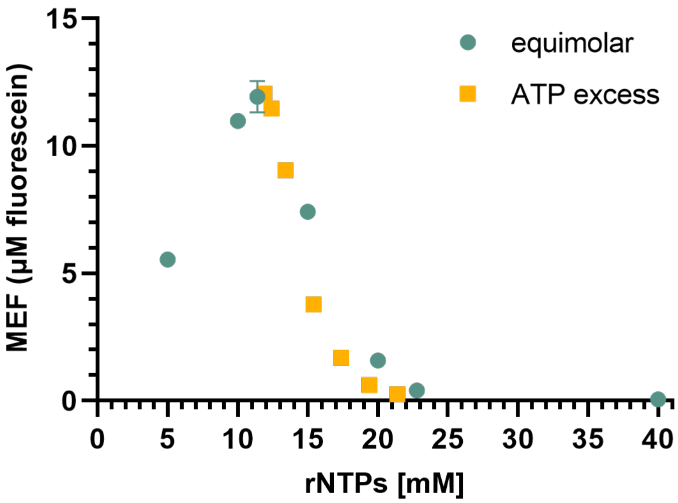


**Supplementary Figure 8:** Titration of rNTPs in constitutive IVT reactions, either in equimolar concentrations (green) or with an excess of ATP (2.85 mM each rNTP + excess ATP). The ATP excess data is the same as shown in Figure 2B-C and Supplementary Figure 7. IVT output with equimolar rNTPs peaks at 11.4 mM rNTPs (2.85 mM each rNTP). Additional nucleotides decrease IVT output due to the chelation of free magnesium ions by additional phosphate groups. Interestingly, the addition of only ATP boosts the decreasing trend, suggesting competition effects. Data are the mean of *n*=3 technical replicates ± s.d. of 4 h time points. All IVT reactions were performed in 40 mM HEPES buffer pH 7.8 and were templated with 15 nM linear GlcR sensor DNA.

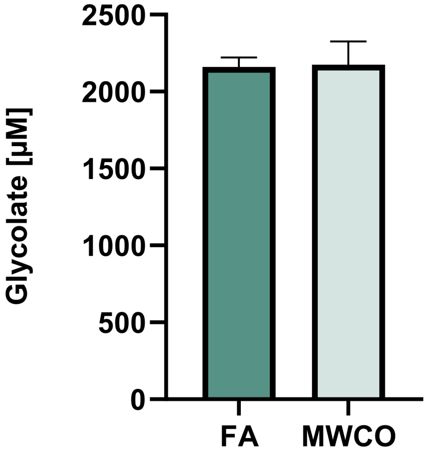


**Supplementary Figure 9:** LC-MS quantification of glycolate production from CETCH cycle samples, either quenched by adding 5% formic acid (FA) or filtered through a 10 kDa MWCO membrane. CETCH cycle composition is the best condition from Pandi et al. (day 7, condition 15[16]). Data are the mean of *n*=2 technical replicates ± s.d.

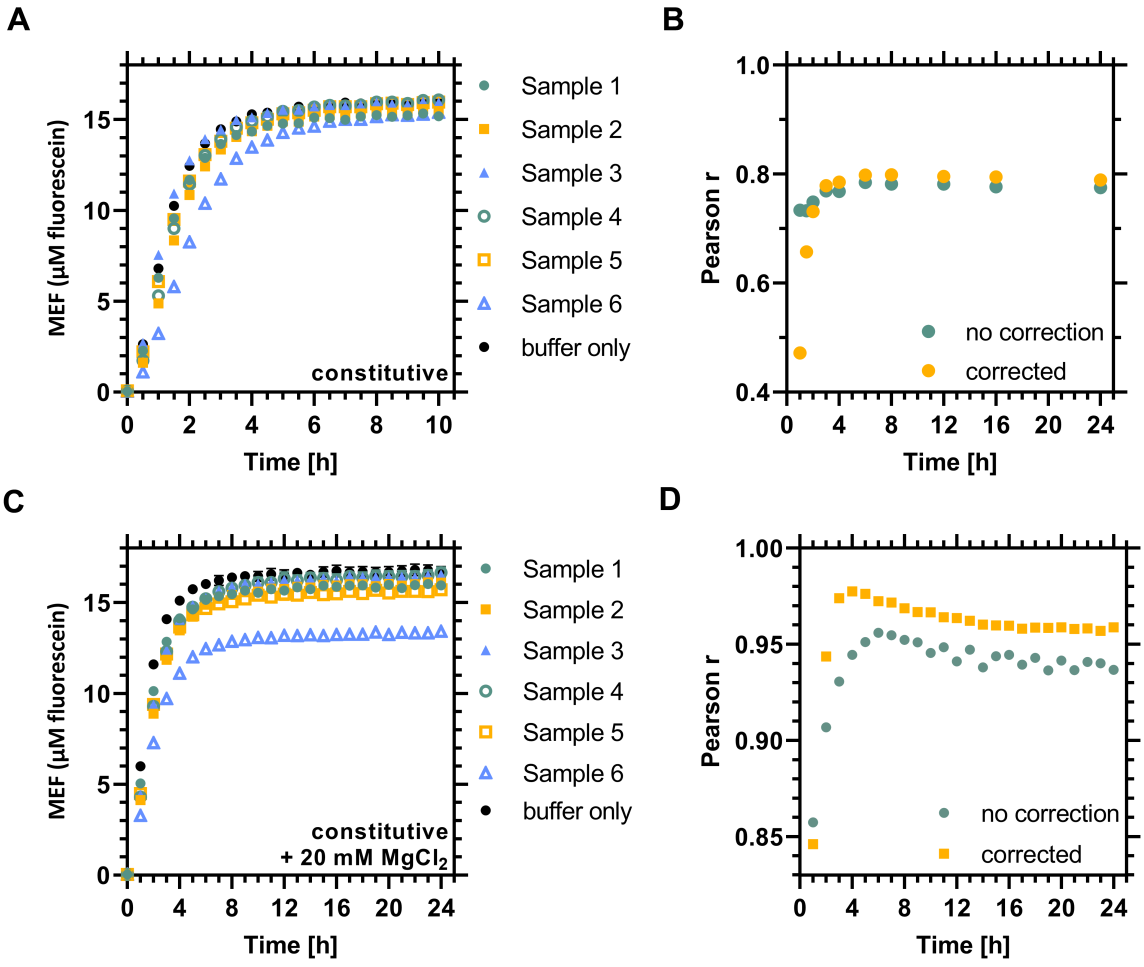


**Supplementary Figure 10:** Effect of CETCH cycle samples on constitutive IVT output. **A:** Constitutive IVT expression in absence of GlcR and with respective CETCH samples added (1:10 diluted in the system). **B:** Correlation of LC-MS quantification and GlcR sensor module output under standard conditions, with and without correction (see description below). **C:** Constitutive IVT expression with additional 20 mM in absence of GlcR and with respective CETCH samples added (1:10 diluted in the system). **D:** Correlation of LC-MS quantification and GlcR sensor output with additional 20 mM $MgCl_2$, with and without correction (see description below). **Applied correction term:** CETCH cycle samples influence the constitutive IVT output, so we decided to apply a correction factor to account for glycolate-independent effects on the IVT system. The measured GlcR sensor output is multiplied by the proportion of the constitutive output of the buffer only control to the constitutive output of the sample at the respective time point ($corrected\ IVT\ output(t) = sensor\ output_{sample}(t) * \frac{constitutive\ output_{control}(t)}{constitutive\ output_{sample}(t)}$). This measure proves helpful to increase the correlation of the IVT output to LC-MS measurement, but also reduces the throughput of an experiment by 2-fold, because samples need to be measured in a constitutive IVT reaction as well. Data are the mean of *n*=3 technical replicates ± s.d. over a 24 h time course. All IVT reactions were performed in 40 mM HEPES buffer pH 7.8 and were templated with 15 nM linear GlcR sensor DNA.

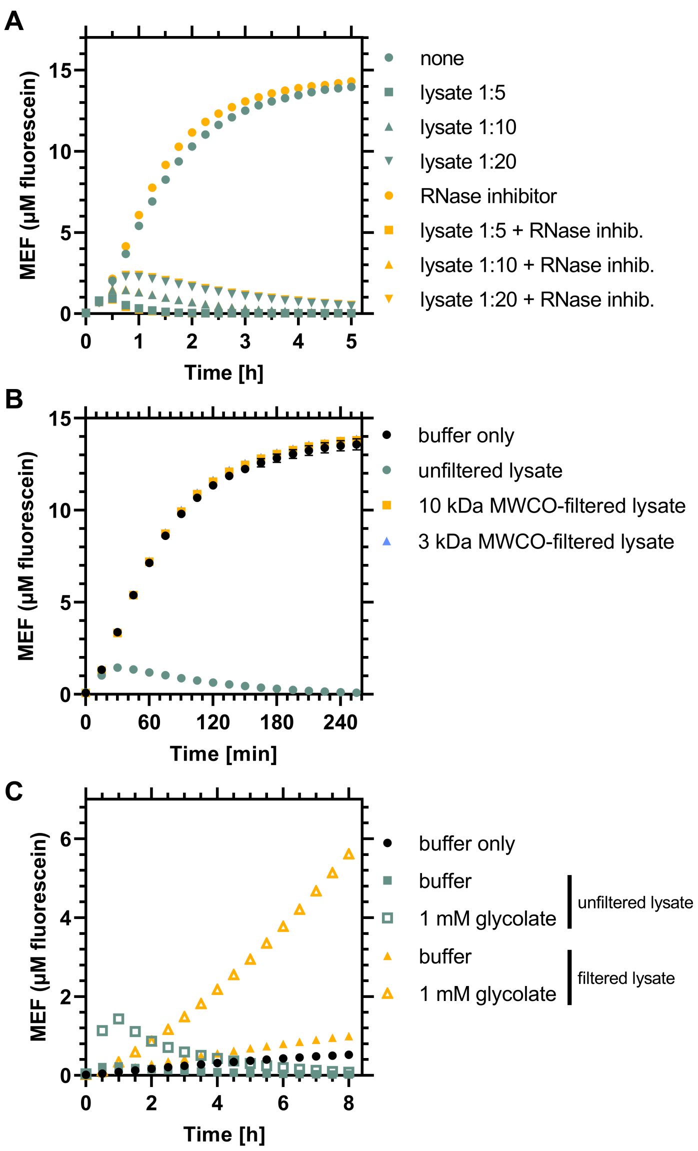


**Supplementary Figure 11:** Testing compatibility of ROSALIND with *E. coli* BL21 Star lysate, and how to avoid RNA degradation. **A:** Titration of lysate in constitutive IVT, either with (orange) or without (green) murine RNase inhibitor (New England Biolabs, 0.5 µL/reaction, # M0314S). The addition of lysate strongly impacts IVT readout in a dose-dependent manner. The decline in fluorescence points to RNA degradation which is not prevented by the added RNase inhibitor. **B:** Testing the effect of 3 kDa or 10 kDa MWCO-filtered lysate samples on constitutive IVT. No significant difference between buffer and filtered lysate samples is visible, while the IVT readout declines in the presence of unfiltered lysate. **C:** Testing the behavior of the GlcR sensor when adding lysate to which either glycolate or buffer was added. Only in filtered lysate samples, the addition of glycolate significantly increase the IVT output. Data are the mean of *n*=3 technical replicates ± s.d. over a 4-8 h time course. All IVT reactions were performed in 40 mM HEPES buffer pH 7.8 and templated with 15 nM linear GlcR sensor DNA. 750 nM MGlcR was added to the experiment in **C**, MGlcR was omitted in the experiments of **A** and **B**.

**Supplementary References**


1. Jung, J. K. *et al.* Cell-free biosensors for rapid detection of water contaminants. *Nature Biotechnology* (2020) doi:10.1038/s41587-020-0571-7.
2. Jung, J. K., Alam, K. K. & Lucks, J. B. ROSALIND: Rapid Detection of Chemical Contaminants with In Vitro Transcription Factor-Based Biosensors. in (eds. Karim, A. S. & Jewett, M. C.) 325–342 (Springer US, New York, NY, 2022). doi:10.1007/978-1-0716-1998-8_20.
3. Jung, J. K., Archuleta, C. M., Alam, K. K. & Lucks, J. B. Programming cell-free biosensors with DNA strand displacement circuits. *Nature Chemical Biology* (2022) doi:10.1038/s41589-021-00962-9.
4. Hanko, E. K. R. *et al.* A genome-wide approach for identification and characterisation of metabolite-inducible systems. *Nature Communications* **11**, 1213 (2020).
5. Hanko, E. K. R., Joosab Noor Mahomed, T. A., Stoney, R. A. & Breitling, R. TFBMiner: A User-Friendly Command Line Tool for the Rapid Mining of Transcription Factor-Based Biosensors. *ACS Synthetic Biology* **12**, 1497–1507 (2023).
6. d'Oelsnitz, S., Love, J. D., Diaz, D. J. & Ellington, A. D. GroovDB: A Database of Ligand-Inducible Transcription Factors. *ACS Synthetic Biology* **11**, 3534–3537 (2022).
7. d'Oelsnitz, S. *et al.* Using fungible biosensors to evolve improved alkaloid biosyntheses. *Nature Chemical Biology* (2022) doi:10.1038/s41589-022-01072-w.
8. d'Oelsnitz, S., Nguyen, V., Alper, H. S. & Ellington, A. D. Evolving a Generalist Biosensor for Bicyclic Monoterpenes. *ACS Synthetic Biology* (2022) doi:10.1021/acssynbio.1c00402.
9. Snoek, T. *et al.* Evolution-guided engineering of small-molecule biosensors. *Nucleic Acids Research* **48**, e3–e3 (2020).
10. Ellefson, J. W., Ledbetter, M. P. & Ellington, A. D. Directed evolution of a synthetic phylogeny of programmable Trp repressors. *Nature Chemical Biology* **14**, 361–367 (2018).

11. Sun, Z. Z., Yeung, E., Hayes, C. A., Noireaux, V. & Murray, R. M. Linear DNA for Rapid Prototyping of Synthetic Biological Circuits in an Escherichia coli Based TX-TL Cell-Free System. *ACS Synthetic Biology* **3**, 387–397 (2014).
12. Lehr, F.-X. *et al.* Modular Golden Gate Assembly of Linear DNA Templates for Cell-free Prototyping. Preprint at https://doi.org/10.48550/arXiv.2310.13665 (2023).
13. Stukenberg, D. *et al.* The Marburg Collection: A Golden Gate DNA Assembly Framework for Synthetic Biology Applications in Vibrio natriegens. *ACS Synthetic Biology* **10**, 1904–1919 (2021).
14. Shimizu, Y. *et al.* Cell-free translation reconstituted with purified components. *Nature Biotechnology* **19**, 751–755 (2001).
15. Schada von Borzyskowski, L. *et al.* Multiple levels of transcriptional regulation control glycolate metabolism in Paracoccus denitrificans. *bioRxiv* 2024.03.11.584432 (2024) doi:10.1101/2024.03.11.584432.
16. Pandi, A. *et al.* A versatile active learning workflow for optimization of genetic and metabolic networks. *Nature Communications* **13**, 3876 (2022).
